# Longitudinal, multi-platform metagenomics yields a high-quality genomic catalog and guides an *in vitro* model for cheese communities

**DOI:** 10.1101/2022.07.01.497845

**Authors:** Christina C. Saak, Emily C. Pierce, Cong B. Dinh, Daniel Portik, Richard Hall, Meredith Ashby, Rachel J. Dutton

## Abstract

Microbiomes are intricately intertwined with human health, geochemical cycles and food production. While many microbiomes of interest are highly complex and experimentally intractable, cheese rind microbiomes have proven powerful model systems for the study of microbial interactions. To provide a more comprehensive view of the genomic potential and temporal dynamics of cheese rind communities, we combine longitudinal, multi-platform metagenomics of three ripening washed-rind cheeses with whole genome sequencing of community isolates. Sequencing-based approaches revealed a highly reproducible microbial succession in each cheese, co-existence of closely related *Psychrobacter* species, and enabled the prediction of plasmid and phage diversity and their host associations. Combined with culture-based approaches, we established a genomic catalog and a paired 16-member *in vitro* washed rind cheese system. The combination of multi-platform metagenomic time-series data and an *in vitro* model provides a rich resource for further investigation of cheese rind microbiomes both computationally and experimentally.

**Importance:** Metagenome sequencing can provide great insights into microbiome composition and function and help researchers develop testable hypotheses. Model microbiomes, such as those composed of cheese rind bacteria and fungi, then allow the testing of these hypotheses in a controlled manner. Here, we first generate an extensive longitudinal metagenomic dataset. This dataset reveals successional dynamics, yields a phyla-spanning bacterial genomic catalog, associates mobile genetic elements with their hosts and provides insights into functional enrichment of *Psychrobacter* in the cheese environment. Next, we show that members of the washed-rind cheese microbiome lend themselves to *in vitro* community reconstruction. This paired metagenomic data and *in vitro* system can thus be used as a platform for generating and testing hypotheses related to the dynamics within, and functions associated with, cheese rind microbiomes.

## Introduction

Microbiomes play crucial roles in human health^1^, geochemical cycles^2^ and food production^3^. While the characterization of community composition of diverse microbiomes has come a long way, our mechanistic understanding of community functioning, and thus our ability to predict and manipulate it, lags behind^4^. Given the complexity and experimental intractability of many microbiomes of interest, model microbiomes consisting of a manageable number of community members that are experimentally tractable can help facilitate the generation and testing of hypotheses about microbiome function. Cheese rind communities have already provided valuable insights into the biology of microbiomes and microbial interactions within them, such as cross-feeding of amino acids and competition for iron; they have also helped uncover the importance of fungi in driving microbial interactions ^5–13^. The cheese rind model communities currently available for *in vitro* studies are based on natural and bloomy rind communities and represent low-complexity microbiomes ranging from 3-7 species ^5, 6, 14, 15^.

In contrast to natural and bloomy rind cheeses, washed-rind cheeses are produced by regular washing (or smearing) with a brine solution^16^. As such, the microbial communities on the surface of washed-rind cheeses experience homogenization throughout its development, which may facilitate intermicrobial interactions or evolutionary processes such as horizontal gene transfer. To date, several studies have examined the community composition of washed-rind cheeses using culture-dependent and culture-independent techniques^5, 17–26^. These studies have shown that bacteria often outnumber fungi by orders of magnitude^19, 20^. Among the bacterial community members, Actinobacteria such as *Brevibacterium, Corynebacterium* and *Glutamicibacter*, and Proteobacteria, such as *Psychrobacter* and *Halomonas,* are usually detected in these communities^17–22^. Lastly, it has been shown that the communities associated with these cheeses show reproducible community succession^19^. Emerging metagenomic techniques have the potential to enable a deeper characterization of the species- and strain-level diversity, functional potential, eco-evolutionary dynamics, and mobile genetic elements within these communities.

In this study, we combine several metagenomics techniques (amplicon, short-read and long-read shotgun sequencing and metaHi-C) with a longitudinal dataset of three washed-rind cheese communities that were collected over the course of cheese ripening. This data provided insights into the reproducible successional trajectories of the studied washed-rind cheeses, provided a catalog of genomes of community members that can be used as references for future studies, identified plasmids and phages and associated them with their host genomes, and provided insights into the biology of these communities. For example, we investigated the striking diversity of *Psychrobacter* genomes recovered from the communities and present a functional enrichment study that highlights enrichment of genes involved in type six secretion as well as siderophore acquisition, two traits that are of high value in the densely populated, iron-limited environment of the cheese rind. Finally, we used the genomic catalog to establish a representative culture collection of washed rind community members and reconstituted an *in vitro* model communities based on the washed-rind cheese microbiome. This model microbiome contains 16 members from several microbial phyla, including several *Psychrobacter* representatives, allowing the examination of interactions across diverse microbial groups and representing the most complex cheese rind-based model microbiome to date. We show that this *in vitro* community undergoes reproducible succession and, by removing certain taxonomic groups, we start to investigate the interaction dynamics in this community.

## Results

### Longitudinal sampling of three washed-rind cheeses from the same facility

Rind samples of three different types of washed-rind cheeses (Cheese A, B, and C) were collected from triplicate batches (each batch made approximately 1 week apart) at six time points throughout ripening (**Fig. 1A**). Cheeses A, B, and C differ with respect to their milk source, use of pasteurized vs raw milk (pasteurized milk was used for cheeses A and B, raw milk is used for cheese C), production location, and production methods. However, during the aging process of each of the cheeses, similar aging practices were used, such as repeated washing with brine solution, and all were aged in the same facility. For all samples, DNA was extracted and 16S and ITS amplicon sequencing were performed. Subsequently, samples from batch 3 were characterized using in-depth metagenomic sequencing, including short-read metagenomic sequencing of all six time points, long-read metagenomic sequencing of rinds at weeks 2, 3, 4, 9 and 13, and metaHi-C of weeks 2, 4, and 13 in the case of Cheeses B and C, and weeks 2 and 13 in the case of Cheese A (**Suppl. Fig. 1, Suppl. Table 1, Suppl. Table 2**).

**Figure 1.**
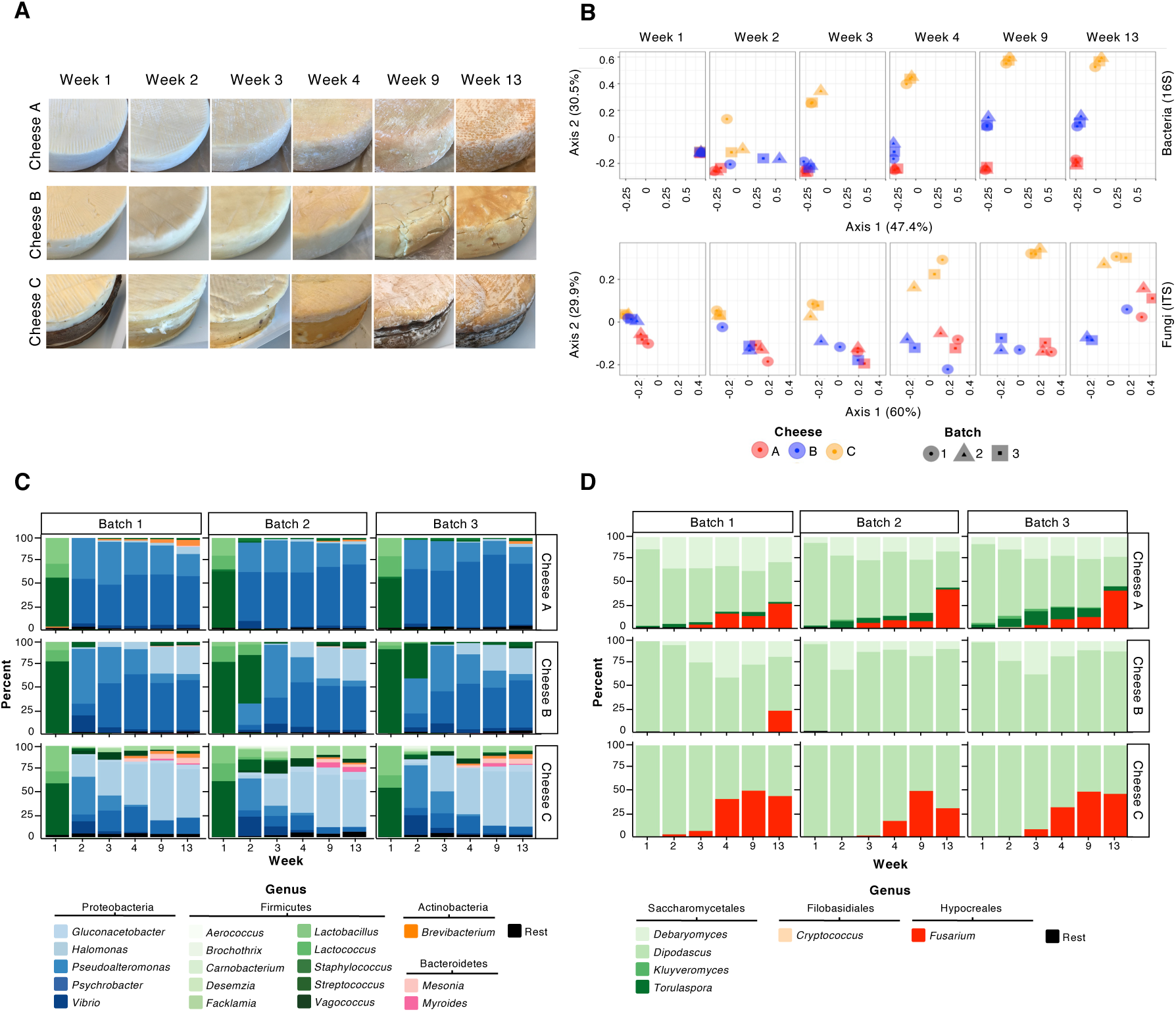
Three washed-rind cheeses from the same facility show reproducible succession patterns during ripening. **(A)** For each of the three different washed-rind cheeses (A, B and C), we followed the aging of three different batches produced one week apart. From each batch we collected rind from duplicate wheels at six time points. A detailed sequencing overview and collection schedule are found in **Suppl.** Fig. 1 and **Suppl. Table 1**, respectively. Representative images of each of the three cheeses at different time points are shown. Cheese C is wrapped in spruce during ripening. In the pictures for weeks 2-4 the spruce has been removed. **(B)** The 16S and ITS data of the ripening communities were subjected to principal component analyses of the Bray-Curtis dissimilarities. Colors correspond to different cheeses and shapes correspond to different batches. **(C, D)** Relative abundance plots of (C) bacteria as determined by 16S amplicon sequencing and (D) fungi as determined by ITS sequencing. Shown are the relative abundances of amplicon sequence variants collapsed at the genus-level. Rest = genera with <1% of the classified reads. Detailed information about 16S and ITS read statistics can be found in **Suppl. Tables 3 and 4**, respectively.

### Successional dynamics of the rind communities throughout ripening

To gain a higher-level overview of community dynamics and reproducibility of succession patterns, we analyzed the relative abundance of bacterial and fungal populations using 16S and ITS sequencing (**Suppl. Tables 3 and 4**). The successional dynamics of each of the three cheeses was remarkably reproducible for both the bacterial and the fungal communities (**Fig. 1C, D**). Principal component analysis based on Bray-Curtis indices indicates that while the different cheeses are highly similar at the earliest sampled time points, they diverge from each other along reproducible trajectories throughout aging (**Fig. 1B**). One exception is one batch of cheese B, which clusters more closely to cheese A at the final fungal sequencing time point. This difference is mainly due to *Fusarium,* which is found in all 3 batches of cheese A and detected in the ITS sequences of that batch of cheese B, but not the other two batches of that same cheese (**Fig. 1D**). Indeed, a wheel from this batch has a visibly different rind than cheese wheels from the other two batches (**Supp.** **Fig. 2**). Although at the genus level the three communities diverge over time, there are consistent successional patterns at the phylum and order levels across cheeses. All three cheeses are dominated by Saccharomycetales throughout ripening, while Hypocreales reproducibly establishes in the communities of cheeses A and C, but not cheese B, over time (**Suppl. Fig. 3B**). Regarding the bacterial communities, all three cheeses are dominated by Firmicutes in Week 1, likely due to the lack of rind resulting in sampling of lactic acid bacteria in the cheese core. Proteobacteria quickly take over and dominate the bacterial communities of all three cheeses by the end of ripening. Cheese A also shows a reproducible establishment of Actinobacteria and Cheese C additionally shows a reproducible establishment of Bacteroidetes in the communities (**Suppl. Fig. 3A**).

**Figure 2.**
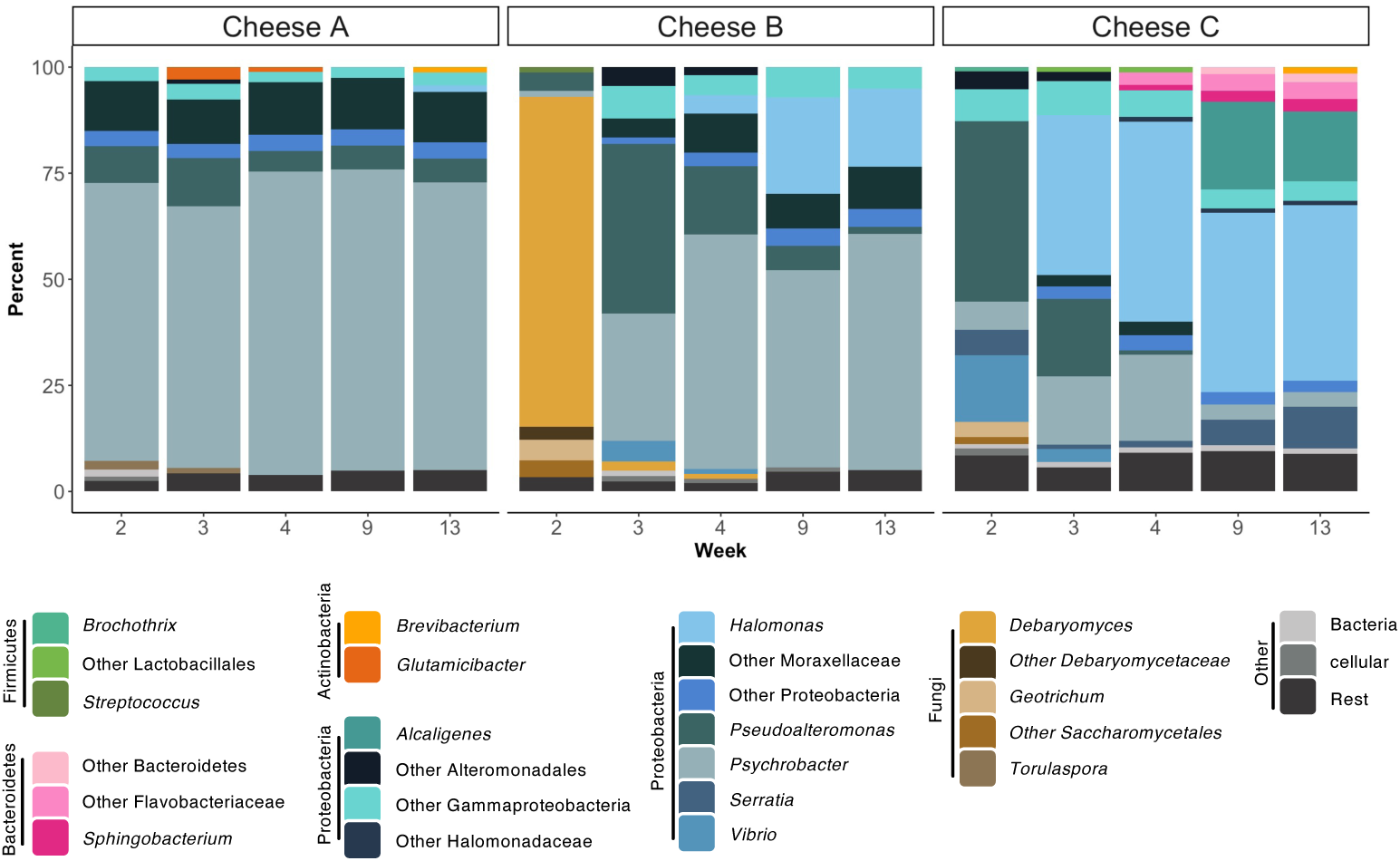
Long-read-based relative abundance reveals dominance of bacterial community members. Relative abundance plots of taxa (aggregated at the genus level or higher) detected by long-read shotgun metagenomic sequencing. Rest = taxa with <1% of the classified reads. Detailed information about read statistics can be found in **Suppl. Table 5.**

While amplicon sequencing provides a high-level overview of community succession and its reproducibility, it does not address the relative abundance of bacteria and fungi to each other. To close this information gap, we performed taxonomic classification of long-read shotgun metagenomic sequencing data (**Suppl. Table 5**). The majority of reads were classified to at least the genus level (**Suppl. Fig. 4).** Long-read-based taxonomic classifications revealed that all three cheeses were heavily dominated by bacteria and that the fungi only constituted a small proportion of the communities, especially at the end of ripening (**Fig. 2, Suppl. Fig. 5).** Consistent with the amplicon sequencing, long-read based taxonomy revealed genera that are shared between all three cheeses, such as *Psychrobacter* and *Pseudoalteromonas*, as well as genera that were specific certain cheeses, such as *Alcaligenes* and *Sphingobacterium*, which are specific to Cheese C (**Fig. 2, Suppl. Fig. 5**).

Overall, Cheese C contained the largest number of unique taxa per rank at all sampled time points (**Suppl. Fig. 6**), which is in accordance with the amplicon results. Cheese C also shows the highest degree of taxa turnover; it is initially dominated by *Pseudoalteromonas*, which is then taken over by *Halomonas* and *Alcaligenes*. In addition, gram-positive community members such as *Brevibacterium* and *Sphingobacterium* become more abundant over time in Cheese C than they do in the other two communities. In contrast, Cheese B is initially dominated by the yeast *Debaryomyces*, before a bacterial community dominated by *Psychrobacter* and *Pseudoalteromonas* takes over. Eventually, *Pseudoalteromonas* is largely displaced from the Cheese B community, while *Halomonas*, members of the family Moraxellaceae, and other unidentified Proteobacteria become more abundant. In contrast to both Cheeses B and C, Cheese A shows very minimal community succession between weeks 2 and 13 and is dominated by *Psychrobacter* and other members of the family Moraxellaceae throughout this entire period of ripening.

### A phyla-spanning genomic catalog of washed-rind cheese communities

We next aimed to generate a genomic catalog for each of the three cheeses to help provide insights into sub-genus diversity, functional potential, and mobile genetic element diversity of the communities. To do so, the long-read shotgun data was assembled for each cheese either by timepoint or across all time points (co-assemblies) (**Suppl. Table 6).** The assemblies were then binned to generate metagenome-assembled genomes (MAGs) (**Suppl. Table 7)**. In addition, community members from Cheese B were isolated and subjected to short- and long-read sequencing for *de novo* hybrid genome assembly (**Suppl. Table 8)**. For each cheese, we combined all high-quality MAGs from the individual timepoint assemblies, the co-assemblies, and the isolate genomes (for cheese B). We then de-replicated each dataset and selected representative MAGs for each cheese. For selection of representative MAGs/genomes from a given dereplication group, isolate genomes were given the highest priority, followed by circular MAGs from individual timepoint assemblies, then circular MAGs from co-assemblies, then complete, non-circular MAGs from individual timepoint assemblies and finally complete, non-circular MAGs from co-assemblies. If several bins per group fell into the same category, the MAGs were prioritized based on quality as assessed by CheckM.

For cheeses A and C, we recovered 17 and 37 high-quality MAGs, respectively (**Fig. 3**, **Suppl. Tables 9 and 11**). In cheese A, 12 out of the 17 high-quality MAGs were both single-contig and circular. In the case of cheese C, 24 of the 37 high-quality MAGs were single-contig and 19 of those were circular. For cheese B, we recovered 11 high-quality MAGs and 16 isolate genomes (**Suppl. Table 10**). Four out of the 11 high-quality MAGs were both single-contig and circular. Consistent with our amplicon sequencing data, comparing the genomic catalogs recovered from the three cheeses using ANI values reveals that the cheeses contain both common and unique genomes (**Suppl. Fig. 7**). Specifically, when considering an ANI cut-off of 99, we observed that six MAGs were represented in the genomic catalogs from all three cheeses (**Fig. 3B**). Based on GTDB-Tk^27^, these MAGs were annotated as *Mesonia* sp., *Vibrio casei, Pseudoalteromonas nigrifaciens, Psychrobacter alimentarius, Vibrio litoralis,* and *Pseudoalteromonas pyrdzensis.* Additionally, cheeses A and B have 8 MAGs (and isolate genomes) in common, cheeses A and C share 2 MAGs and cheeses B and C share 5 MAGs (and isolate genomes) (**Suppl. Table 12**). Lastly, cheese A has 1 unique MAG, cheese B has 8 unique MAGs (and isolate genomes) and cheese C has 24 unique MAGs.

**Figure 3.**
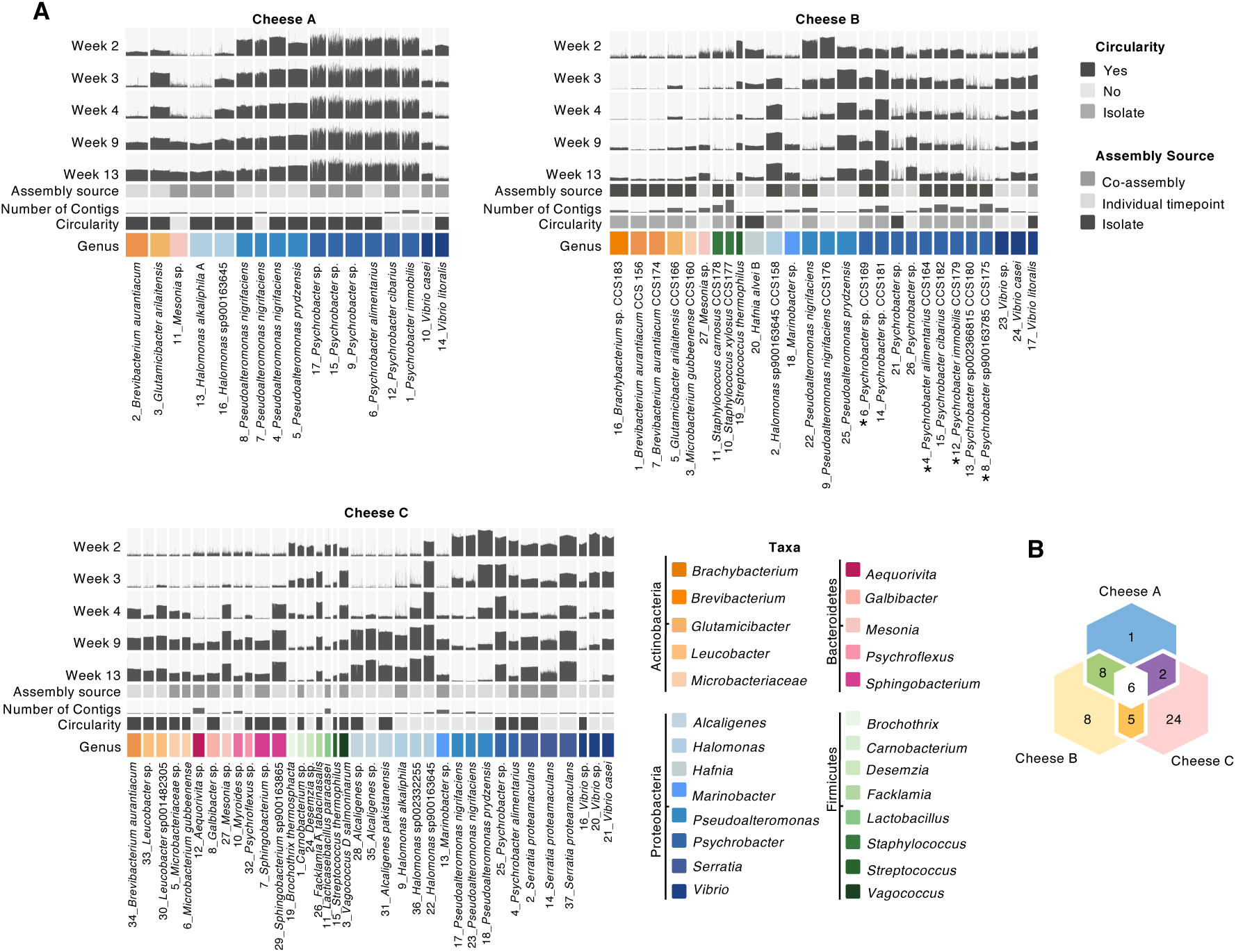
A phyla-spanning bacterial genomic catalog of three washed-rind cheeses. **(A)** Anvi’o plots show dereplicated high-quality metagenome-assembled genomes (MAGs) (and isolate genomes in the case of cheese B). MAGs were generated either by assembling HiFi-reads from individual time points or by co-assembling reads from all time points of a cheese. For each MAG, the number of contigs is indicated (lowest value being 1, highest value being 22). Circularity of MAGs is indicated as is the taxonomy as predicted by GTDB-Tk. Colors correspond to genera. The bin names contain the genomic catalog bin number and the predicted taxonomy of the bin. To estimate relative abundances of MAGs over time, the short-reads from weeks 2, 3, 4, 9 and 13 were mapped to the genomic catalog for each cheese. **Suppl. Table 7** shows all non-dereplicated bins. Genomes marked with a * were included in *in vitro* community reconstruction. **(B)** Overlap of high-quality MAGs (and isolate genomes) from cheeses A,B, and C, with MAGs having an ANI >99% considered to be the same.

Altogether, the genomic catalog covers the bacterial phyla Proteobacteria, Firmicutes, Bacteroidetes and Actinobacteria. For each cheese, Proteobacteria are the group with the most MAGs (and isolate genomes) with 14, 18, and 18 MAGs (and isolate genomes) representing this phylum in Cheeses A, B and C, respectively. Further, we are able to recover multiple species and/or strain representatives for several genera. For Cheeses A and B we observe multiple co-occurring *Psychrobacter* species and/or strains. Six out of 14 and nine out of 18 Proteobacteria are of the genus *Psychrobacter* for Cheeses A and B, respectively. Both cheeses also contain a diversity of distinct *Pseudoalteromonas* representatives with four and three distinct MAGs (and isolate genomes) recovered from Cheeses A and B, respectively. Cheese C also contains two and three distinct MAGs belonging to the genera *Psychrobacter* and *Pseudoalteromonas*, respectively. In addition, we observed species and/or strain-level diversity in the genera *Alcaligenes, Brevibacterium, Halomonas, Leucobacter, Microbacterium, Serratia, Sphingobacterium, Staphylococcus* and *Vibrio.* We further note that four of 17, eight of 27 and 15 of 37 MAGs (and isolate genomes) from Cheeses A, B, and C respectively, were not classified at the species level by GTDB-Tk, which could indicate potentially new species in our genomic catalog.

To gain an overview of how abundant each of the recovered MAGs (and isolate genomes) was during community development, the communities were also subjected to short-read metagenomic sequencing, and the reads were mapped to the genomes in the genomic catalog (**Fig. 3**). For almost all time points, more than 50 percent of the reads were mapped to the genomic catalog (**Suppl. Table 13**). The only exception is Cheese B at week two, in which only about 19 percent of the reads mapped to the genomic catalog. From the long-read based community composition analysis (**Fig. 2**), we know that at this time point Cheese B is dominated by a fungus. Since the genomic catalog provided in **Fig. 3** only contains bacterial genomes, it is not surprising that a low amount of short reads from week two maps against the bacterial genomic catalog for Cheese B. In contrast, for some time points over 90 percent of the short reads from Cheeses B and C map to the genomic catalog, indicating that the catalog provides a good cross-section of the bacterial communities at these time points.

The availability of strain-resolved genomic catalogs next allowed us to interrogate the successional dynamics of each cheese more deeply. For example, amplicon and long read-based taxonomy suggested that there were relatively small changes in community composition over time in Cheese A. However, short read mapping back to the genomic catalog revealed species and strain-level temporal dynamics. Specifically, *Pseudoalteromonas nigrifaciens* initially slightly dominates over *Pseudoalteromonas prydzensis* in Cheese A, while by the end of ripening in week 13 *Pseudoalteromonas prydzensis* slightly dominates over *Pseudoalteromonas nigrifaciens*. Similarly, *Vibrio casei* initially dominates over *Vibrio litoralis* in Cheese A, while already by week three, *Vibrio litoralis* starts to dominate over *Vibrio casei.* Very similar dynamics are also observed in Cheese B for *Pseudoalteromonas* and *Vibrio*. *Pseudoalteromonas nigrifaciens* initially dominates over *Pseudoalteromonas prydzensis* before finally being taken over by it. Similarly, *Vibrio litoralis* initially dominates over *Vibrio casei* before *Vibrio casei* catches up in terms of relative abundance towards the end of ripening. For both Cheeses A and B, we saw little variation in abundances across the *Psychrobacter* MAGs (and isolate genomes), potentially due to a high similarity between the MAGs (and isolate genomes) or to stable coexistence of species/strains over time. For Cheese C, the abundance of *Pseudoalteromonas* MAGs decreases over time, however their relative abundance to each other does not change significantly. For Psychrobacter, only one of the two representative MAGs decreases in abundance over time, while the other shows a stable abundance based on read mapping.

### Meta-HiC and long reads associate viruses and plasmids with their hosts

In addition to generating a genomic catalog for each cheese, we were interested in leveraging the existing long-read metagenomics data to characterize the diversity of plasmids and viruses within these microbiomes. To help capture any short and/or low abundance mobile genetic elements that might have been missed in the long-read assemblies, we first generated mega-assemblies of each cheese combining both long-read and high-depth short-read datasets. In brief, the short metagenomic shotgun reads were mapped to the co- and individual timepoint assemblies and any reads that did not map were assembled. The resulting short read-based contigs were combined with the de-replicated unbinned long read-based contigs and the contigs from the de-replicated MAGs to yield mega-assemblies for each of the three cheeses. These mega-assemblies were subjected to plasmid prediction and virus prediction (**Suppl. Table 14, Suppl. Fig. 8**). We identified 419, 297 and 343 putative plasmid contigs in Cheeses A, B and C, respectively. Additionally, we identified 109, 116 and 214 lysogenic and 40, 32 and 67 lytic virus contigs for Cheeses A, B and C, respectively. Of those, four, four, and seven contigs were classified as complete, circular viruses, while 22, 32 and 57 contigs were classified as high quality draft viruses for Cheeses A, B and C, respectively. We last considered the distributions of the predicted viruses (**Suppl. Fig. 9**) and plasmids (**Suppl. Fig. 10)**. In this regard, Cheese C stands out for the presence of several predicted viruses with sizes larger than 200,000 basepairs, which is a hallmark of jumbophages^28^.

We next used the Hi-C data generated for various time points to identify putative hosts for the extrachromosomal MGEs (plasmids and lytic phages). MGE predictions were used as inputs for the viralAssociationPipeline^29, 30^ program together with the total mega-assemblies, the metaHi-C reads, and the long reads (**Fig. 4, Suppl. Fig. 11, Suppl. Table 15)**. Altogether, 70, 21 and 120 of the predicted extrachromosomal MGEs associate with MAGs, while an additional 209, 105 and 89 extrachromosomal MGEs associate with non-MAG (unbinned) contigs for cheeses A, B and C, respectively. One hundred seventy-three, 199 and 195 of the extrachromosomal contigs remain unassociated (**Suppl. Fig. 11)**. Of the extrachromosomal MGEs associated with MAGs, we identify 40, 11 and 96 MGEs for Cheeses A, B and C, respectively, that associate with a MAGs (or isolate genome) at one timepoint only. Thirty-one, 10, and 23 MGEs from Cheeses A, B and C, respectively, associate with the same MAG (or isolate genome) at two time points and one MGE each for Cheese B and Cheese C associate with the same MAG at three time points. The vast majority of these putative MGEs was not associated with a putative host through the initial binning. For Cheeses A, B and C, seven, one, and 11 MGEs, respectively, associate with more than one MAG. One MGE, a predicted plasmid (s673.ctg001008l_6, indicated by an asterisk in **Fig. 4**), associates with two different MAGs at two different time points, indicating a putative HGT event. At week two, this MGE is associated with a *Psychrobacter* sp. host and at week 13, this same MGE is associated with a *Pseudoalteromonas prydzensis* host. Interestingly, this plasmid is predicted to encode genes involved in iron uptake, which have previously been associated with horizontally-transferred regions in cheese ^31^. The full results of the association pipeline are found in **Suppl. Table 15.**

**Figure 4.**
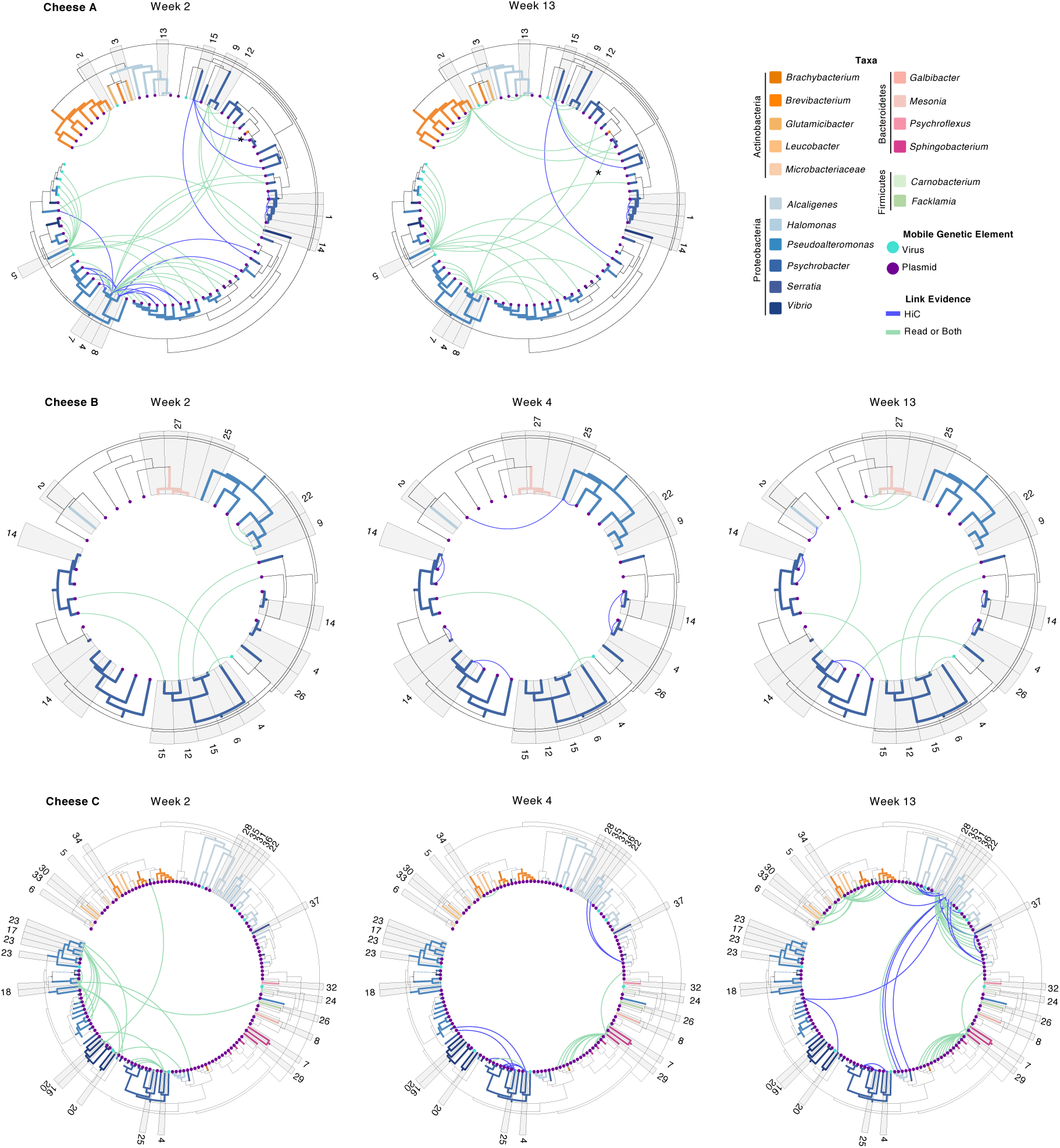
metaHi-C and long read-based evidence of MGE associations with hosts from the washed-rind cheese genomic catalog. iTOL graphs show the long read-based or metaHiC-based association of MGEs (lytic viruses and plasmids) with MAGs (and isolate genomes) from the genomic catalog. The branches of the trees represent contigs and are colored by the contigs’ predicted taxa. Contigs that are classified as lytic viruses and plasmids are indicated with colored dots (teal = virus, purple = plasmid). Bin numbers for MAG host contigs are shown on the outside of the tree. Associations between MGEs and their hosts are indicated by the lines. Blue lines indicate metaHiC-based evidence for association, while green lines indicate long read-based evidence or both metaHiC and long read-based evidence. The asterisk indicates the associations for contig s673.ctg001008l_6. The full results from the viralAssociationPipeline.py program can be found in **Suppl. Table 15.**

We next looked at MGEs that changed their genus association over time (**Suppl. Table 16**). Twelve, six, and three MGEs from Cheeses A, B and C, respectively, changed hosts over the course of sampling based on taxonomic prediction of the host contig. We identified several predicted instances of interkingdom host changes, especially in Cheese B between *Geotrichum* and proteobacterial species. Again, we identified iron uptake genes associated with potentially transferred elements. Some of these elements also contain genes related to phosphonate transport, which has also previously been observed in association with horizontally-transferred iron uptake regions in cheese^31^.

### Pangenome analysis of *Psychrobacter* and functional enrichment in cheese isolates

Our genomic catalog revealed high sub-genus level diversity within the *Psychrobacter* genus. To investigate conserved and unique gene sets across *Psychrobacter* species, we aimed to analyze the *Psychrobacter* pangenome. We first determined the non-redundant *Psychrobacter* isolate genomes and MAGs from the combined three-cheese dataset. This resulted in a total of seven isolate genomes and two MAGs. An additional 97 publicly available *Psychrobacter* genomes were combined with these nine genomes in a pangenomic analysis (**Fig. 5A, Suppl. Table 17**). The 97 publicly available genomes were sourced from marine, soil, host-associated, cheese, other fermented food, and other miscellaneous environments . We attempted to include at least one representative of every *Psychrobacter* species possessing a publicly available genome. As expected, pangenomic analysis identified a core set of genes common to *Psychrobacter* from diverse environments. This core gene set, defined as present in 95% of genomes, consisted of 1045 genes and made up 9.4 percent of total gene clusters, with the remaining 90.6 percent of gene clusters classified as accessory by panX (**Fig. 5A**). A core-genome SNP tree constructed from all variable positions of all single copy core genes showed some evidence of clustering of genomes by environment and, with the exception of *Psychrobacter immobilis*, by species (**Fig. 5B**). These data suggest that *Psychrobacter* species may have environment-specific gene sets. Functional enrichment analysis of gene clusters was used to find functions (clusters of orthologous groups (COGs)) that were enriched in genomes of cheese isolates compared to genomes from other environments. Specifically, a group of genes related to iron access, particularly through the use of iron-chelating siderophores, was enriched in *Psychrobacter* genomes from cheese (adj. q-value < 0.1, **Fig. 5C, Suppl. Table 18**). In addition, genes related to type VI secretion systems, contractile defense systems that bacteria can use to transport effector proteins into target cells, were also enriched in cheese *Psychrobacter* genomes relative to other environments (adj. q-value < 0.1, **Fig. 5C**). An alignment of type VI secretion gene regions identified in the cheese genomes revealed a consistent organization in genomes from cheeses made in geographically separated origins. sEleven of the 13 cheese genomes contained two distinct type VI clusters in separate genomic regions, one larger cluster (often with *impA* to *impH*) and one smaller cluster (often with *impI* to *impM*) (**Fig. 5D**). The larger region was not identified in *Psychrobacter* sp. str. FME6. In *Psychrobacter* sp. str. CCS181, these two regions are adjacent to each other in the genome. These type VI clusters were often bounded by tRNA/tmRNA regions. In two strains, CCS179 and JB193, these clusters also contain phage-related gene sets within the tRNA/tmRNA boundary points in addition to the type VI genes. Both of these clusters were predicted by PHASTER to contain intact prophage^32, 33^.

**Figure 5.**
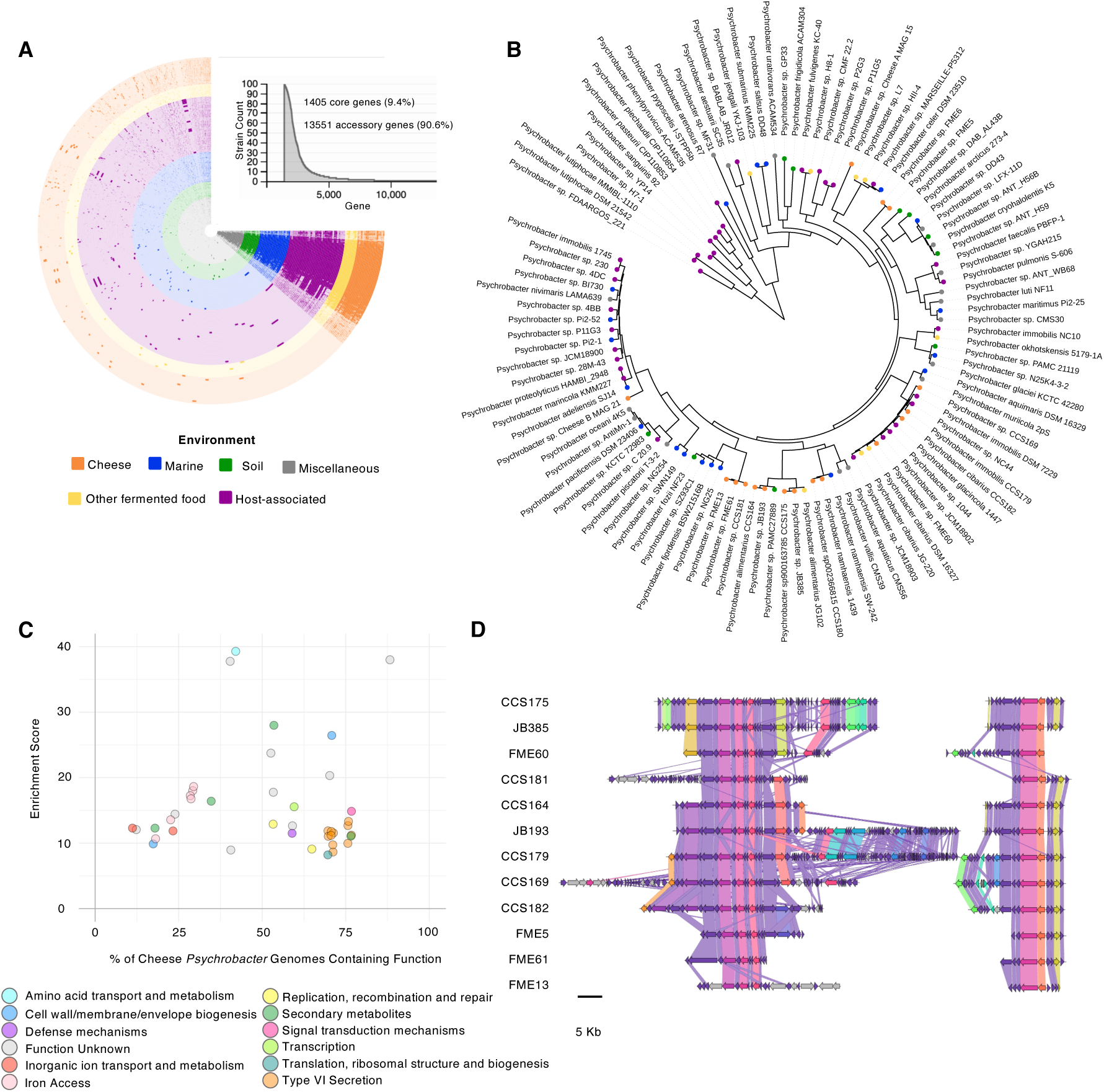
Pangenomic analysis of *Psychrobacter* genomes from diverse environments. **(A)** Pangenomic analysis was performed on 106 genomes from marine, soil, cheese, other fermented food, and miscellaneous other environments. **(B)** Phylogenetic tree of *Psychrobacter* genomes based on alignment of single copy core genes. **(C)** Functional enrichment of COG categories in *Psychrobacter* genomes from cheese relative to other environments. **(D)** Alignment of the two type VI secretion gene clusters from *Psychrobacter* genomes from various cheeses.

### *In vitro* community reconstruction

Given the utility of previous model systems based on cheese rind microbiomes, we next aimed to establish whether the washed-rind cheese communities lend themselves to *in vitro* experimentation. To this end we selected 16 microbial species (13 isolates from Cheese B, three from previous isolation efforts) that would represent both the breadth and depth of diversity found in a typical washed-rind cheese microbiome. Given that the metagenomic analysis presented here revealed that multiple species and strains of the genus *Psychrobacter* can co-occur, we chose to include 4 *Psychrobacter* isolates from Cheese B (*Psychrobacter alimentarius* CCS164, *Psychrobacter sp.* CCS169, *Psychrobacter sp*900163785 CCS175 and *Psychrobacter immobilis* CCS179). These isolates represent two different groups of *Psychrobacter* (**Fig. 5B**). In addition to the *Psychrobacter* isolates, the community also contains four other Proteobacteria: *Pseudoalteromonas nigrifaciens* CCS176*, Halomonas sp900163645* CCS158*, Vibrio casei* JB196*, and Hafnia alvei* JB232; two Firmicutes: *Staphylococcus xylosus* CCS177 and *S. carnosus* CCS178; four Actinobacteria: *Brevibacterium linens* JB5, *Brachybacterium ssp.* CCS183, *Glutamicibacter arilaitensis* CCS165, and *Microbacterium gubbeenense* CCS160; and two fungi: *Debaryomyces hansenii* CCS145 and *Galactomyces geotrichum* CCS187. The selected cheese rind isolates were then combined into *in vitro* communities following established protocols. In brief, a total of 100,000 CFUs per community member were inoculated on the surface of a 10% Cheese Curd Agar petri dish^14^. To mimic the production process for washed-rind cheeses, the surfaces of the plates were washed with 20 percent NaCl using a sterile swab every 24 hours in the first 96 hours for a total of four washes. Communities were incubated in the dark at 15℃ in a humidified environment (**Suppl. Fig. 12)**. To assess the reproducibility of the *in vitro* washed cheese rind model, CFU counts of each sampling day (days three, five, seven, 21) were done for both the bacterial and fungal members (**Suppl. Fig 13, Suppl. Table 19**). In addition, a portion of each sample from sampling days seven and 21 was used for DNA extraction and short-read metagenomic sequencing. Sequencing reads were then mapped back to reference genomes of the bacterial community members to track their relative abundance (**Fig. 6A, Suppl. Fig. 14, Suppl. Table 20**). The additional sequencing and mapping allowed for analysis of *Psychrobacter* at the species level and revealed that all four species persist within the full community. However, the relative proportion of each species varied over time, with *Psychrobacter CCS169* most abundant at day 7, and the other three strains, especially *Psychrobacter alimentarius CCS164,* increasing in relative abundance at day 21 (**Fig. 6A**).

**Figure 6.**
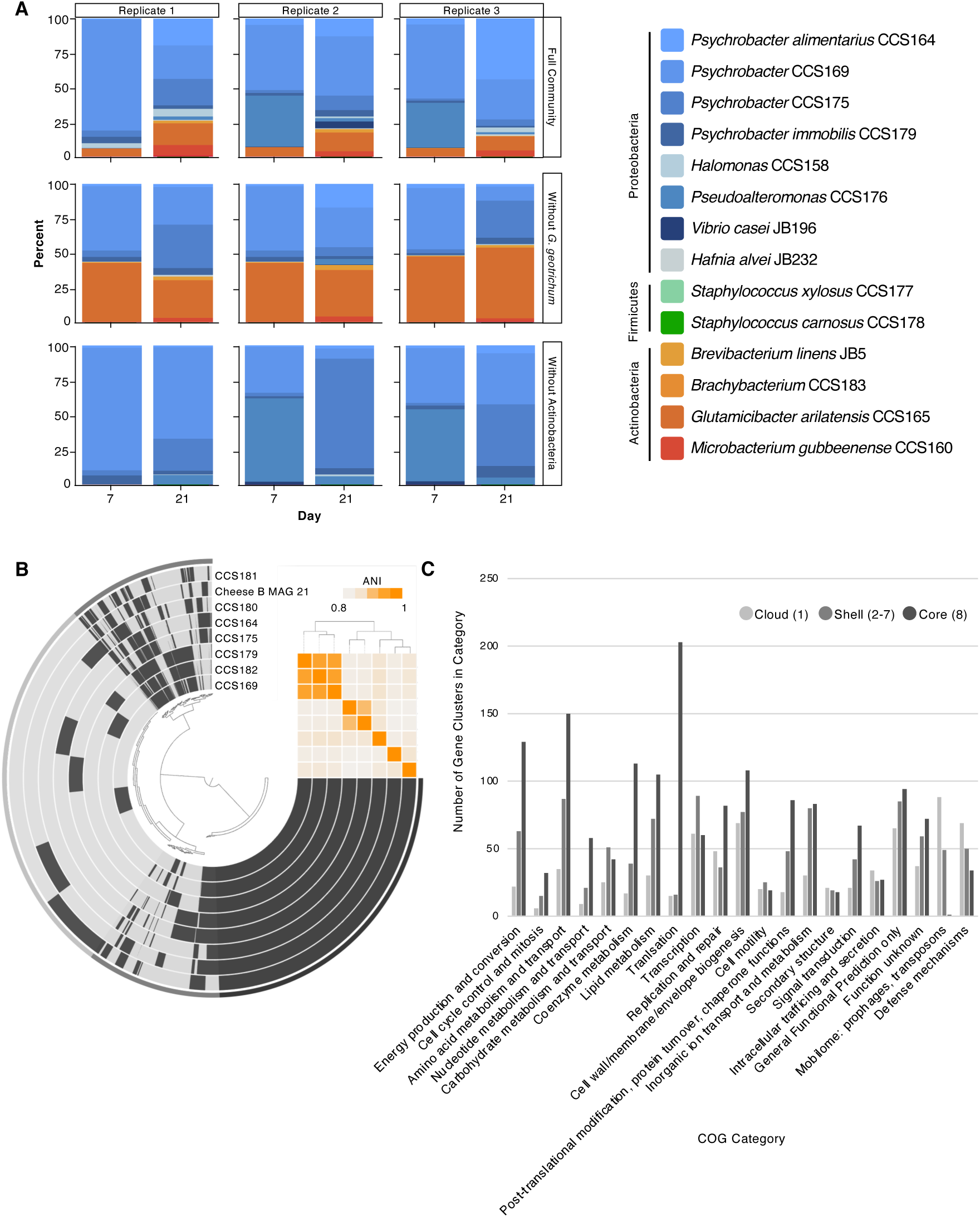
A 16-member *in vitro* model system based on the washed-rind cheese microbiome. **(A)** Relative abundance plots of the *in vitro* community after seven and 21 days of growth based on short read mapping to reference genomes. **(B)** Pangenomic analysis of eight *Psychrobacter* genomes from Cheese B. **(C)** Functional categories of core (present in all eight genomes), shell (in two-seven genomes), and cloud (unique to one genome) gene clusters from eight *Psychrobacter* genomes from Cheese B.

Previous studies revealed a positive correlation between *G. geotrichum* and gamma-proteobacterial species, and a negative correlation between *G. candidum* and actinobacterial species based on co-occurrence patterns in a sequencing based survey of cheese rind microbiomes^5^. In addition, pairwise experimental data showed stimulatory effects of *G. geotrichum* on gamma-proteobacterial growth and inhibitory effects on actinobacterial growth^5^. We took advantage of the fact that this model contains all three of these members (*G. geotrichum*, Proteobacteria, and Actinobacteria) to examine whether these patterns hold in a community context. To do this, we reconstructed a community lacking *G. geotrichum* and evaluated whether this resulted in any differences in the final community composition. The removal of *G. geotrichum* favored the relative growth of the Actinobacteria over the Proteobacteria, even though overall bacterial absolute abundance was very similar between these two samples (**Fig. 5A**). Absolute abundance based on read counts reveals that the decrease in gamma-proteobacterial abundance is largely due to the poor growth of *Pseudoalteromonas* (**Suppl. Table 19)**. In contrast, almost all of the actinobacterial species reach higher absolute abundances compared to the full community. Overall, these results are consistent with *G. geotrichum* inhibiting Actinobacteria and stimulating Proteobacteria.

We next tested the effect of removing all actinobacterial members from the community. Similar to the other conditions, *Psychrobacter* species dominate the communities, but increase to 93-95% in the community without Actinobacteria. These communities cluster separately from the full community and community without *G. geotrichum* in principal component analysis (**Suppl. Fig. 15**). In contrast to the other communities, *Pyschrobacter alimentarius CCS164* does not increase in relative abundance, remaining below 5% of *Psychrobacter* in all replicates. Instead *Psychrobacter CCS169* competes with *Psychrobacter CCS175* for the most abundant *Psychrobacter*.

To investigate the genetic basis of the coexistence of multiple *Psychrobacter* species within a single cheese, which was supported by our sequencing and *in vitro* growth data, we did an additional pangenomic analysis focused specifically on the eight *Psychrobacter* genomes from Cheese B (MAG and isolate genomes) (**Fig. 6B**). This analysis identified a set of ‘cloud’ gene clusters unique to each genome, ‘shell’ gene clusters found in at least 2 genomes but not all genomes, and ‘core’ gene clusters found in all eight genomes. As accessory gene sets may provide clues to the coexistence of multiple closely-related species or strains, we looked at the COG functional categories associated with the core, shell, and cloud gene sets. While central metabolic processes like amino acid metabolism and translation were most prevalent in the core set, the cloud set was enriched in defense mechanisms and the mobilome (phages and transposons)(**Fig. 6C, Suppl. Table 21**). As these gene categories may be involved in interspecies interactions, this may be an interesting starting point for future experiments to investigate the mechanisms underlying species and/or strain co-occurrence.

## Discussion

Washed rind cheeses harbor unique, moderate complexity microbiomes that may provide useful systems for the study of microbial succession, interactions, strain-level dynamics and horizontal gene transfer. This work represents a comprehensive, in-depth analysis of a set of washed rind cheese microbiomes combining culture-independent and culture-dependent approaches. Using amplicon and metagenomic sequencing, we showed that the three washed-rind cheeses that were the subject of this study show reproducible succession patterns. Both amplicon sequencing and long-read taxonomic classification show that the mature communities of the investigated cheeses are dominated by bacteria, in particular Proteobacteria. Overall, we detect very similar bacterial and fungal genera as previous studies investigating the composition of washed-rind cheeses^5, 17–26^, but provide new insights into species and strain-level diversity and dynamics.

We also provide a catalog of high-quality MAGs (and isolate genomes) that spans the bacterial phyla Firmicutes, Bacteroidetes, Actinobacteria, and Proteobacteria. In fact, two of the three cheeses yielded representative genomes from all four of those phyla despite low proportional abundance. It is likely that the long read sequencing helped not only recover those low abundance genomes, but also helped in the resolution of species and/or strain-level MAGs. For example, we were able to resolve several *Psychrobacter* genomes in the three cheeses which may have been difficult to resolve using short read sequencing alone.

We utilized the bacterial genomic catalog generated as part of this study to further investigate the biology within these microbiomes. First, we leveraged the long reads together with metaHi-C reads to associate putative mobile genetic elements with their hosts. Many of these mobile genetic elements were not binned with their putative hosts during the initial binning and only became associated in this additional analysis. We were able to assign 15, six, and 29% of extrachromosomal MGEs to MAG hosts and 46, 32, and 22% of extrachromosomal MGEs to non-MAG contigs for Cheeses A, B, and C, respectively. Using this method we also detected several putative HGT events, including mobile genetic elements predicted to encode iron-uptake pathways. Previous examination of cheese bacterial genomes revealed a strong enrichment of iron-uptake pathways on horizontally-transferred regions^31^. The changes in host association as suggested by metaHi-C in this study require further *in vitro* confirmation and follow-up to fully understand their exact nature and impact within the cheese microbiome.

Next, we investigated the striking diversity of *Psychrobacter* within the communities and investigated functional enrichment within cheese-associated *Psychrobacter* MAGs and isolate genomes as compared to *Psychrobacter* genomes associated with other environments. We identify an enrichment of iron acquisition genes as well as type VI secretion genes in cheese-associated *Psychrobacter*. Both of these functional enrichments are reasonable given what is known about the cheese rind environment. First, the cheese microbiome has evolved to thrive in iron-limiting conditions^8, 34, 35^. Second, the dense microbial communities that make up cheese rind microbiomes may lead to contact-dependent microbial interactions. Type VI secretion systems enable the delivery of cargo proteins that modulate bacterial-bacterial and bacterial-eukaryotic interactions^36^. As such, contact-dependent interaction machinery enriched in cheese-associated *Psychrobacter* may play a larger role in species interactions in the densely populated cheese microbiome as opposed to interactions relevant to free-living, marine *Psychrobacter*, for example. Future *in vitro* studies could aim to understand how type VI secretion-mediated dynamics contribute to community composition and function.

The comprehensive, longitudinal nature of this dataset makes it a potentially valuable resource for future investigations into the biology of cheese microbiomes or the study of microbiome dynamics in general. For example, the approaches applied here could be used to interrogate other types of MGEs, such as integrative and conjugative elements and prophages. Of particular interest in this dataset would be the >300,000 bp prophage predicted in cheese C, which might be a putative jumbophage. Additionally, one aspect of the long read data, which we did not explore in this paper, is the fact that it can detect methylation patterns on DNA, which can help associate extrachromosomal elements with their hosts and could theoretically be used to identify HGT events^37^. Indeed, a recent study used this approach to recover MAGs from a marine microbial consortium and identify HGT events, phage infection and strain-level structural variation^38^.

Finally, we showed that, similar to previous work on cheese microbiomes^5, 6^, washed rind communities are amenable to *in vitro* reconstruction. The *in vitro* system established here represents a higher level of diversity than previous models, and includes species and strain-level diversity. The pairing of an experimental microbiome system with a corresponding in-depth metagenomic dataset should provide ample future opportunities for generating and testing hypotheses related to the many facets of biology present within these microbiomes.

## Materials and Methods

### Cheese rind sample collection and DNA extraction

Three production batches of three different washed-rind cheeses were sampled over the course of ripening starting from fresh cheese wheels. At weeks 1, 2, 3, 4, 8 and 12 of ripening the cheesemakers overnight shipped two wheels of each sampled batch. The cheeses were stored at 4 ℃ for up to 48 h prior to harvest. During the harvest, rind was scraped off each wheel using fresh razor blades as in Wolfe *et al*., 2014^5^. The rind samples were homogenized by gentle stirring with sterile pipette tips. A rice grain-sized portion of these samples was frozen in cryotubes at -80 ℃ in phosphate-buffered saline + 40% glycerol. Where applicable, 10 mg of each wheel were collected in sterile 2-ml tubes for cross-linking as part of the ProxiMeta Hi-C library preparation (see below). Finally, 500 mg of each sample were set aside in sterile 1.5-ml tubes at 4 ℃ for DNA extraction. DNA was extracted within 72h using a phenol-chloroform extraction protocol. If DNA was extracted at a later time, the rind samples were stored at -80 ℃.

For the DNA extraction, the rind samples were ground into powder in liquid nitrogen (https://lab.loman.net/2018/05/25/dna-extraction-book-chapter/#RefHeading_Toc505877552). The rind powder was incubated for 1 hour at 37 ℃ in 7 ml modified Tris-lysis buffer (10 mM Tris-Cl (pH 8), 100 mM EDTA (pH 8), 1% SDS, 20 µg/ml RNAse A, 20 mg/ml lysozyme). 87.5 µl of Proteinase K (800 units/ml) (New England Biolabs, Ipswich, MA, USA) were added, the samples were mixed by inversion and incubated at 50 ℃ for 1 hour. The samples were then subjected to two rounds of DNA extraction using equal volumes of phenol-chloroform isoamyl alcohol. The final aqueous phase was mixed with equal volumes of ice cold isopropyl and 0.1 volume of 3M sodium acetate. The precipitated DNA was pelleted in 5 ml centrifuge tubes at 17,000 g for 3 minutes. When the volume of the final aqueous phase was larger than 5 ml, the supernatant was removed and the remaining sample was added to the same centrifuge tubes to repeat the pelleting step. The final pellets were washed in 1 ml ice cold 70% ethanol and air-dried for about 15 min. Last, the DNA was resuspended in 500 µl UltraPure™ DNase/RNase-Free Distilled Water (ThermoFisher Scientific, Waltham, MA, USA) and kept at room temperature overnight to allow the DNA to dissolve. DNA was then stored at - 20 ℃ until further processing.

### 16S and ITS amplicon sequencing and analysis

For the 16S and ITS sequencing of the community DNA samples, we followed the Illumina-supplied “16S Metagenomic sequencing Library Preparation” protocol (Part # 15044223 Rev. B) and “Fungal Metagenomic Sequencing Demonstrated Protocol” (Document # 1000000064940 v01), respectively. The 16S-specific primers (16S forward: 5’-TCGTCGGCAGCGTCAGATGTGTATAAGAGACAG-CCTACGGGNGGCWGCAG-3’, 16S reverse: 5’-GTCTCGTGGGCTCGGAGATGTGTATAAGAGACAG-GACTACHVGGGTATCTAATCC-3’) targeted the 16S V3 and V4 regions. The ITS-specific primers (ITS forward: 5’-TCGTCGGCAGCGTCAGATGTGTATAAGAGACAG-CTTGGTCATTTAGAGGAAGTAA-3’,ITS reverse: 5’-GTCTCGTGGGCTCGGAGATGTGTATAAGAGACAG-GCTGCGTTCTTCATCGATGC-3’) were the same as the ITS_fwd_1 and ITS_rev_1 primers listed in the “Fungal Metagenomic Sequencing Demonstrated Protocol”. For each of the two amplicon PCRs, 1.25 µl DNA (5 ng/µl) from the duplicate wheels for each sample were mixed and the resulting 2.5 µl were amplified either with the 16S- or the ITS-specific primers and Q5^®^ Hot Start High-Fidelity DNA Polymerase (New England Biolabs, Ipswich, MA, USA). The PCR settings used were as follows: initial denaturation at 98 ℃ for 3 minutes, 25 rounds of: 98 ℃ - 10 seconds, 55 ℃ - 30 seconds, 72 ℃ - 15 seconds, final extension at 72 ℃ for 2 minutes.

Amplicon PCRs were purified using AMPure XP beads (Beckman Coulter, Indianapolis, Indiana, USA) and eluted swith UltraPure™ DNase/RNase-Free Distilled Water (ThermoFisher Scientific, Waltham, MA, USA). For the index PCR, the 16S and ITS amplicon PCRs were subjected to amplification with IDT for Illumina Nextera DNA Unique Dsual Indexes (Set A, now called IDT for Illumina DNA/RNA UD Indexes) (produced by: Integrated DNA Technologies, Coralville, Iowa, USA, sold by: Illumina, Inc., San Diego, California, USA) and Q5^®^ Hot Start High-Fidelity DNA Polymerase (New England Biolabs, Ipswich, MA, USA).

Index PCRs were again purified using AMPure XP beads (Beckman Coulter, Indianapolis, Indiana, USA) and eluted with UltraPure™ DNase/RNase-Free Distilled Water (ThermoFisher Scientific, Waltham, MA, USA). The final Index PCRs were quantified using the Qubit dsDNA HS kit (ThermoFisher Scientific, Waltham, MA, USA) with the Qubit 2.0 fluorometer (ThermoFisher Scientific, Waltham, MA, USA), diluted to 2 nM and pooled at equimolar ratios. The pools were diluted to the final loading concentration of 50 pM and spiked with 50 pM PhiX control v3 (Illumina, Inc., San Diego, California, USA) to a final concentration of 10% PhiX. 20 ul of the 16S(+PhiX) and of the ITS(+PhiX) pools were sequenced individually in-house on an iSeq 100 (paired-end, 150 bp) (Illumina, Inc., San Diego, California, USA).

The fastq files of the forward reads of the respective sequencing runs were imported into Qiime2 (version 2020.11.1) ^39^ and denoised with Dada2 ^39–41^ (qiime dada2 denoise-single --i- demultiplexed-seqs with flags --p-trim-left 0 --p-trunc-len 0 --p-n-threads 16). The denoised forward ITS reads were classified using classify-sklearn with a Naive Bayes classifier (qiime feature-classifier fit-classifier-naive-bayes) trained on a custom ITS database^5, 39, 40, 42–44^. The denoised forward 16S reads were classified in the same way using a pre-trained Greengenes database (Greengenes 13_8 99% OTUs full-length sequences (MD5: 03078d15b265f3d2d73ce97661e370b1))^39, 40, 42–46^. Detailed read statistics can be found in Suppl. **Tables 1 and 2**. Ordination plots (method: PCoA, distance: Bray-Curtis) were generated using R (version 4.0.2) (https://www.r-project.org/) together with the qiime2R (version 0.99.4), tidyverse (version 1.3.1) and phyloseq (version 1.34.0) packages^47–49^ . The stacked bar plots were also created with R using the qiime2R, tidyverse and phyloseq packages. Taxonomies were collapsed at the genus level for both the ITS and the 16S data. For the stacked bar plots showing a higher taxonomic level, taxonomies were collapsed at phylum (16S) or order (ITS) level. In all cases, taxonomies with less than 1% abundance were aggregated under the category “Rest”.

### metaHi-C

10-15 mg of rind from duplicate wheels for each sample were fixed separately and then combined for further processing according to the ProxiMeta kit methodology from Phase Genomics (Seattle, WA). The multiplexed libraries were sequenced by Novogene (Sacramento, CA, USA) on a HiSeq 4000 with a run configuration of 2 x 150 bp.

### Metagenomic shotgun sequencing

For the SMRT sequencing, DNA from duplicate cheese wheels of each sample was combined in equal ratios. 5 µg DNA of each sample were used as input for the library preparation, which was carried out by SNPSaurus (Eugene, Oregon, USA). The DNA was sheared to a modal size of 10 kb using a Megaruptor 2 (Diagenode Diagnostics, Seraing (Ougrée), Belgium). Libraries were prepared using Pacific Bioscience’s Express Template Prep kit version 2.0 according to the manufacturer’s protocol (https://www.pacb.com/wp-content/uploads/Procedure-Checklist-Preparing-Multiplexed-Microbial-Libraries-Using-SMRTbell-Express-Template-Prep-Kit-2.0.pdf, Menlo Park, CA, USA). Samples were pooled two at a time and size selected using a BluePippin (Sage Sciences, Beverly, MA, USA) with the 0.75% Agarose Dye-free 10-18kb cassette, U1 marker and a 10kb+ cutoff. The final libraries were sequenced by the Genomics & Cell Characterization Core Facility (University of Oregon, Eugene, Oregon, USA) on a Sequel II according to the SMRT Link Set Up (SMRT cell type = 8M, Sequencing kit = v2.0, Sequencing primer = v2, Binding kit = v2.0, Sequencing control = v1, Polymerase binding time = 4 hr, Movie time = 30 hr, Pre-extension time = 2 hr, Loading concentration = 100, 133, 150, or 200 pM (varied by cell), Loading method = Diffusion). HiFi reads were generated using the CCS tool (https://ccs.how/) version 6.2.0 with default settings (ccs in.subreads.bam out.subreads.bam \ --min-passes 3 \ --min-snr 2.5 \ --min-length 10 \ --max-length 50000 \ --min-rq $MIN_RQ, $MIN_RQ is set accordingly: ccs.Q20 - $MIN_RQ = 0.99, ccs.Q30 - $MIN_RQ = 0.999, ccs.Q40 - $MIN_RQ = 0.9999).

Short-read sequencing libraries were prepared using the Nextera DNA Flex Library Prep workflow (Illumina, Inc., San Diego, California, USA). For each sample, input DNA consisted of DNA from two duplicate cheese wheels from the same sample that was mixed in equal proportions (ng). For multiplexing, IDT for Illumina Nextera DNA Unique Dual Indexes (Set A, now called IDT for Illumina DNA/RNA UD Indexes) (produced by: Integrated DNA Technologies, Coralville, Iowa, USA, sold by: Illumina, Inc., San Diego, California, USA) were used. Pooled libraries were sequenced by the IGM Genomics Center at the University of California San Diego (San Diego, CA, USA) on the NovaSeq 6000 System using both lanes of a NovaSeq SP flow cell and a run configuration of 2 x 250 bp.

### Long-read based relative abundance estimations of washed-rind cheese communities

To investigate successional dynamics of the cheese communities based on HiFi reads we utilized the Pacific Biosciences supplied toolkit “PB-metagenomics-tools” (https://github.com/PacificBiosciences/pb-metagenomics-tools). First, HiFi reads were taxonomically classified with the “Taxonomic-Profiling-Nucleotide” pipeline, which uses minimap2^50^ to align sequences to the NCBI nt database and MEGAN-LR^51^ to interpret alignments and assign reads to taxa. The read counts by taxonomy were generated using the “MEGAN-RMA summary”. Outputs from the “MEGAN-RMA summary” were further processed in R (version 4.0.2) (https://www.r-project.org/) together with the janitor (version 2.1.0), tidyverse (version 1.3.1) and cowplot (version 1.1.1) packages to generate the stacked bar plot in **Fig. 2**. Additionally, the outputs from the “MEGAN-RMA summary’’ were used as inputs for the “compare-kreport-taxonomic-profiles” script available in the PB-metagenomics-tools suite to generate **Suppl. Figs. 4-6**.

### Generation of metagenome-assembled genomes

Metagenomic assemblies were performed with hifiasm-meta (v. 0.2-r053)^52^ using the default settings. For the individual timepoint assemblies, reads were input separately by time point and cheese. For the co-assemblies, reads from all time points were combined by cheese. To evaluate the assemblies and identify high-quality MAGs, the PacBio HiFi-MAG-Pipeline was used (https://github.com/PacificBiosciences/pb-metagenomics-tools, part of the “PB-metagenomics-tools” suite). This pipeline uses minimap2^50^ to align HiFi reads to the contigs to obtain coverage estimates, which are used with MetaBat2^53^ to perform binning using all contigs. A separate bin set is also constructed from all circular contigs (e.g., one bin per circular contig), and the two binning strategies are compared and merged using DAS_Tool^54^. The dereplicated bins are evaluated using CheckM^55^, and quality thresholds are applied to retain high-quality MAGs (defaults of >70% completeness, <10% contamination, <20 contigs). The high-quality MAGs are then analyzed using the Genome Taxonomy Database Toolkit (GTDB-Tk)^27^, which attempts to identify the closest reference genome and assign taxonomy for each MAG.

### Isolation of bacterial community members

The glycerol stocks of one wheel of Cheese B at weeks 2 and 13 were thawed on ice, homogenized by vortexing and pipetting and aliquoted into smaller working stocks, which were frozen at -80 ℃ for at least 24 h before further processing. A dilution series of one working stock each of weeks 2 and 13 was plated on plate count agar supplemented with milk and salt (PCAMS, 5 g/L tryptone, 2.5 g/L yeast extract, 1g/L dextrose, 1 g/L whole milk powder, 10 g/L sodium chloride and 15 g/L agar) containing 100 µg/ml cycloheximide to select against the fungal community members. The plates were kept in a 15 ℃ incubator for 4 days and then in the light at room temperature for an additional 3 days to allow for pigment formation. Colonies with as many distinct colony morphologies as could be distinguished by eye were then purified by re-streaking three times on plain PCAMS, each time with an incubation at 15 ℃ for 4 days and an additional 3-day incubation in the light at room temperature. Colonies from the final re-streak were patched onto plain PCAMS and incubated at 15 ℃ for 4 days. 2 ml overnight cultures (LB Miller broth) were inoculated from the patches and grown shaking at 240 rpm at 22 ℃. After 24h, glycerol stocks were prepared using PBS + 20% glycerol and flash-frozen and stored at -80 ℃. The only exception to this was CCS196, which was grown on PCAMS agar at 22 ℃ for 24 h, harvested in PBS+20% glycerol, flash-frozen and stored at -80 ℃.

### Short-read, whole genome sequencing of Cheese B isolates

Isolates (CCS156, 158, 160, 164, 165, 166, 169, 174, 175, 176, 177, 178, 179, 180, 181, 182, 183, 184, 196) were grown from glycerol stocks on plain PCAMS for 2 days at 15 ℃ and an additional 3 (all except CCS183) or 5 days (only CCS183) at room temperature on the benchtop. DNA was extracted following the NexteraTM DNA Flex Microbial Colony Extraction protocol (Document #1000000035294v01) with the following modifications: AMPure XP beads (Beckman Coulter, Indianapolis, Indiana, USA) were used instead of SPB beads and 2 ml cryotubes filled with approx. 250 µl acid-washed beads (1:1 ratio of 425-600 µm and 150-212 µm beads) were used instead of PowerBead Tubes. In addition, cells were collected from the primary streak using a sterile pipette tip. The extracted DNA was quantified using the Qubit dsDNA HS kit (ThermoFisher Scientific, Waltham, MA, USA) with the Qubit 2.0 fluorometer (ThermoFisher Scientific, Waltham, MA, USA) and stored at -20 ℃. Steps 25 - 27 were skipped. Due to low amounts of cellular input, steps 7 and 8 were skipped for CCS183 and instead 50 µl were mixed with 20 µl AMPure beads. For the library preparation a maximum of 500 ng of DNA was mixed with UltraPure™ DNase/RNase-Free Distilled Water (ThermoFisher Scientific, Waltham, MA, USA) for a total volume of 30 µl. The libraries were prepared using the Illumina DNA Prep protocol (Document # 1000000025416 v09) and the IDT for Illumina DNA/RNA UD Indexes (Set A, produced by: Integrated DNA Technologies, Coralville, Iowa, USA, sold by: Illumina, Inc., San Diego, California, USA). Tagmented DNA was amplified for 5 cycles, except CCS191, which was amplified for 8 cycles due to low input amounts. Libraries were diluted to 30 nM using buffer RSB from the Illumina DNA Prep protocol, pooled at equimolar ratios and sequenced by Novogene (Sacramento, CA, USA) on a HiSeq 4000 with a run configuration of 2 x 150 bp.

### Long-read whole genome sequencing of Cheese B isolates

A subset of isolates (CCS156, 158, 160, 164, 165, 166, 169, 174, 176, 179, 180, 181, 182, 183, 184 and 196) was additionally sequenced using Nanopore technology (Oxford Nanopore Technologies, Oxford, United Kingdom) to improve isolate assemblies. Isolates were either grown shaking at room temperature in 2 ml LB Miller broth for at least 24 hours and until culture appeared cloudy (CC156, 158, 160, 164, 165, 166, 169, 174, 176, 179, 180, 182, 183, 184) or they were grown on PCAMS agar plates at room temperature for ∼48 h (CCS181 and CCS196). Cells from liquid cultures were harvested by centrifugation at 10,000 rpm for 5 min, removal of supernatant by pipetting and freezing at -80 ℃. Cells from the agar plates were recovered by adding 2.5 ml PBS onto the plate and dislodging cells with sterile cell scrapers. 1 ml of recovered cell suspension was centrifuged at 10,000 rpm for 5 min and frozen at -80 ℃ after supernatant removal. Cell pellets were thawed and DNA was extracted using the phenol-chloroform protocol detailed above without the liquid nitrogen grinding step (CCS156, CCS165, CCS166 and CCS176) or using the Qiagen Genomic-tip 20/g kit (Qiagen, Venlo, Netherlands) according to manufacturer instruction. DNA of CCS156, CCS165, CCS166 and CCS184 was extracted from the pellet resulting from the full 2 ml culture, while DNA from the rest of the isolates was extracted from pellets resulting from 1 ml of the overnight culture. DNA extractions were quantified using the Qubit dsDNA HS kit (ThermoFisher Scientific, Waltham, MA, USA) with the Qubit 2.0 fluorometer (ThermoFisher Scientific, Waltham, MA, USA) and their quality was assessed with the Tapestation gDNA assay (Agilent, Santa Clara, CA, USA). DNA was stored at -20 ℃ until the library preparation was carried out using the Nanopore kit SQK-LSK110 (the DNA control strand was included in the library preparation for CCS156, CCS165, CCS166 and CCS176, but not the others). Libraries were either sequenced right away using a flongle flow cell (FLO-FLG001) or stored at -80 ℃ until sequencing. For most samples, base calling was done in real time during the sequencing run (CCS156, CCS158, CCS160, CCS164, CCS166, CCS169, CCS174, CCS179, CCS180, CCS181, CCS182, CCS183, CCS184, CCS196) using the fast basecalling model in the MinKnow software (version 21.02.1 running Guppy version 4.3.4 for CCS156, CCS158, CCS166, CCS180, CCS181, CCS183 and CCS196 or version 21.06.0 running Guppy version 5.0.11 for CCS160, CCS164, CCS169, CCS174, CCS179, CCS182 and CCS184) with a minimum qscore of 7. For a couple of samples (CCS165 and CCS176) base calling was done post sequencing using Guppy (version 5.0.16) on the command line with the fast base calling model (config file dna_r9.4.1_450bps_fast.cfg) and the minimum qscore set to 7. We recovered read N50s between 4 - 30 kb. In all cases, the fastq files in the “pass” folders were concatenated by isolate and used for further analysis.

### Isolate genome assemblies

To generate the isolate assemblies, the Illumina reads were first trimmed using Trimmomatic^56^ (version 0.39) using the paired-end mode and arguments ILLUMINACLIP:NexteraPE-PE.fa:2:40:15:2:True SLIDINGWINDOW:4:20 MINLEN:20. The paired and unpaired, trimmed reads of CCS175, CCS177 and CCS178 were assembled using SPAdes^57^ (version v3.13.0, with arguments -t 14 -m 50 -k 33,55,77,99,127). For CCS156, CCS158, CCS160, CCS164, CCS165, CCS166, CCS169, CCS174, CCS176, CCS179, CCS180, CCS181, CCS182, CCS183, CCS184 and CCS196 hybrid assemblies were performed with SPAdes^58^ using both the paired and unpaired, trimmed Illumina reads as well as the Nanopore reads as input (version v3.13.0 with arguments -t 14 -m 50 -k 33,55,77,99,127). Contigs below 1000 bp were removed.

### Generation of genomic catalog

MAGs resulting from co-assemblies and individual timepoint assemblies (and isolate assemblies in case of cheese B) were de-replicated with dRep (version 3.2.2)^59^. CheckM (version 1.1.3)^55^ results were provided as an Excel sheet. Genomes for the genomic catalog were selected by hand from the resulting MASH clusters by prioritizing isolates (for Cheese B) over circular individual timepoint MAGs over circular co-assembly MAGs over complete, non-circular individual timepoint MAGs over complete, non-circular co-assembly MAGs. If several genomes per cluster fell into the same category, the genomes were prioritized based on low contamination and high completeness as determined by CheckM. Taxonomies of the final genomes in the genomic catalog were determined by GTDB-Tk (version 1.6.0, classify workflow)^27^. To determine the abundance of the genomes from the genomic catalog over time, the genomes in the catalog were indexed using bwa index and the Illumina reads from week 2 - 13 were mapped to the this indexed genomic catalog using bwa mem (Version: 0.7.17-r1188)^60^. Before mapping, the short-read metagenomic reads were concatenated by samples across the two lanes of the sequencing run and trimmed using Trimmomatic^56^ (version 0.39) using the paired-end mode and arguments ILLUMINACLIP:NexteraPE-PE.fa:2:40:15:2:True SLIDINGWINDOW:4:20 MINLEN:20. Samtools^61^ (version 1.9 using htslib 1.9) was used to convert .sam files into .bam files with samtools view and .bam files from all time points of each cheese were combined using samtools cat. Samtools flagstat was used to determine the percentage of reads mapped to the genomic catalog. Anvi’o (version 7.1 ‘hope’)^62^ was used to generate the plots in **Fig. 3**.

### Generation of mega-assemblies

For each cheese, contigs from the co- and individual timepoint assemblies were concatenated and contigs <1000 bp were removed. The concatenated assemblies were indexed using bwa index, and short-read metagenomic reads were mapped to this indexed, concatenated assembly using bwa mem (bwa version 0.7.17-r1188)^60^. Before mapping, the short-read metagenomic reads were concatenated by samples across the two lanes of the sequencing run and trimmed using Trimmomatic^56^ (version 0.39) using the paired-end mode and arguments ILLUMINACLIP:NexteraPE-PE.fa:2:40:15:2:True SLIDINGWINDOW:4:20 MINLEN:20. Samtools^61^ (version 1.9 using htslib 1.9) was used to convert .sam files into .bam files with samtools view and .bam files from all time points of each cheese were combined using samtools cat. Samtools view was then used to extract all short-read metagenomic reads that did not map to the indexed, concatenated assemblies. The resulting bam files were converted into fastq files using the bamToFastq command (version 0.5.3) from the BEDTools suite^63^ and fastq_pair from the fastq-pair package (version 1.0)^64^ was used to synchronize the newly generated fastq files. The reads in the resulting fastq files were assembled using SPAdes^58^ using both the paired and unpaired, trimmed Illumina reads as input (version v3.13.0 with arguments -t 14 -m 50 -k 33,55,77,99,127). To generate the mega-assembly for each cheese, unbinned contigs from the co- and individual timepoint assemblies of the respective cheese were first de-replicated. To de-replicate these contigs, a custom nucleotide blast database was created consisting of the unbinned contigs. Unbinned contigs were then compared to this database using blastn^65, 66^ (version 2.10.1+, options: -outfmt "6 qseqid sseqid pident qcovs length qlen slen evalue score" -evalue 1e-6 -perc_identity 99 -word_size 20 -num_threads 12). A contig was considered redundant if there was an overlap of contigs of at least 90%; the smaller contig in this case was considered redundant. Seqkit^67^ (version 2.2.0) was then used to extract the non-redundant unbinned contigs from the full set of unbinned contigs. These non-redundant unbinned contigs were then combined with the contigs resulting from the assembly of the unmapped Illumina reads and the contigs from the selected de-replicated MAGs in the genomic catalog. This mega-assembly should incorporate all non-redundant assembly information captured from the combination of Illumina and PacBio sequencing.

### Identification of mobile genetic elements

Plasmid contigs were identified in each cheese metagenome by analyzing the mega-assemblies (contigs >5000 bp) with viralVerify (https://github.com/ablab/viralVerify, version 1.1) using the -p flag and Pfam database version Pfam33.1. Viruses were identified with VIBRANT^68^ (version 1.2.1). Numbers of predicted plasmid and virus contigs were plotted using R (version 4.0.2) (https://www.r-project.org/) and the readxl (version 1.3.1) and ggplot2 (version 3.3.5).

### Assigning mobile genetic elements to hosts

To assign mobile genetic elements to hosts we used the viralAssociationPipeline.py script (https://github.com/njdbickhart/RumenLongReadASM/blob/master/viralAssociationPipeline.py)2 ^9, 30^. In brief, the contigs in the mega-assemblies for each cheese were classified taxonomically using Kraken2 and a custom Kraken database containing the default Kraken database supplemented with genomes of cheese-associated microbes. The outputs were reformatted in Excel to fit the input requirements for the viralAssociationPipeline.py script. metaHi-C reads were then aligned to the indexed mega-assemblies using bwa mem (Version: 0.7.17-r1188)^60^ (bwa mem -v 1 -t 16 -5SP {mega-assembly} {forward_Hi-C} {reverse_Hi-C}). Reads that mapped to multiple locations in the assemblies were removed using grep -v -e ’XA:Z:’ -e ’SA:Z:’. The resulting .sam files were then converted into .bam files using samtools view^61^. seqtk subseq (https://github.com/lh3/seqtk) was used to extract the predicted MGE contigs as a .fasta file. The lengths of the contigs in this file were determined using samtools faidx^61^ (version 1.9 using htslib 1.9). The viralAssociationPipeline was then run using the -a, -g, -b, -i, -v, -o, -s, -m, and -l flags. iTOL^69^ was used to generate the diagram in **Fig. 4** showing the connections between MAGs (and isolate genomes in the case of cheese B) and MGEs (plasmids as predicted by viral verify -p and lytic phages as predicted by Vibrant). A summary graph showing the numbers of extrachromosomal MGEs associated with MAGs, with unbinned contigs as well as the numbers of unassociated extrachromosomal MGEs was plotted using R (version 4.0.2) (https://www.r-project.org/) and the readxl (version 1.3.1) and ggplot2 (version 3.3.5).

### *Psychrobacter* pangenome analysis

To identify a non-redundant set of *Psychrobacter* genomes from our cheese isolate assemblies and full set of MAGs from the three cheeses, genomes classified by GTDB-Tk (version 1.6.0, classify workflow)^27^ as belonging to the *Psychrobacter* genus were dereplicated with dRep (version 3.2.2)^59^. CheckM (version 1.1.3)^55^ results were provided as an Excel sheet. Genomes were selected by hand from the resulting MASH clusters by prioritizing isolates over circular individual timepoint MAGs over circular co-assembly MAGs over complete, non-circular individual timepoint MAGs over complete, non-circular co-assembly MAGs. This resulted in the selection of two MAGs (Cheese B MAG 21 and Cheese A MAG 15) and seven isolate genomes. An additional 97 publicly available *Psychrobacter* genomes were downloaded from NCBI (https://www.ncbi.nlm.nih.gov/assembly/) or from the JGI Integrated Microbial Genomes & Microbiomes(IMG/M) system (**Suppl. Table 17**). The full data set consisted of 17 genomes from cheese, eight genomes from other fermented foods, 35 host-associated genomes, ten genomes from soil, 19 genomes from marine environments, and 17 genomes from other miscellaneous environments. These 106 genomes were analyzed using the microbial pangenomics workflow in Anvi’o ^62, 70^ (version 7.1 ‘hope’). Specifically, assemblies were run through gene prediction with Prodigal^71^ (version 2.6.3), hits to bacterial single-copy gene collections were identified using HMMER (http://hmmer.org/), and genes were annotated with functions from the NCBI’s Clusters of Orthologous Groups^72^. Identification of core and accessory gene sets and the construction of the phylogenetic tree based on single copy core gene SNPs was done by panX^73^ (version 1.6.0) using FastTree 2^74^ and RaxML^75^. The core gene set was defined here as present in at least 95 percent of genomes. Functional enrichment analysis was performed in Anvi’o with a comparison of genomes from cheese versus those not from cheese, a corrected q-value cutoff of 0.1, and COG20_FUNCTION as the annotation source^76^. Regions associated with type VI secretion were extracted from cheese isolate genomes manually in Geneious Prime (www.geneious.com) and aligned and visualized using clinker and clustermap.js with default settings^77^. The separate pangenomic analysis of the 8 *Psychrobacter* genomes from cheese B was performed as described for the full data set, and the anvi-summarize function in Anvi’o was used to determine the core, shell, and cloud gene sets and associated COG functional categories. The summary chart was produced based on the Anvi’o output using Microsoft Excel for Mac Version 16.53.

### *In vitro* washed-rind community reconstitution

For the *in vitro* community reconstitution, 16 cheese isolates were chosen based on the metagenomic analyses. From this study, four *Psychrobacter* strains (CCS164, CCS169, CCS175 and CCS179), one *Halomonas* strain (CCS158), one *Pseudoalteromonas* strain (CCS176), one *Glutamicibacter* strain (CCS165), two *Staphylococcus* strains (CCS177, CCS178), one *Brachybacterium* strain (CCS183), one *Microbacterium* strain (CCS160) and two fungal strains (CCS145 - *Debaryomyces hansenii* and CCS187 - *Galactomyces geotrichum*) were chosen. In addition, three strains from a previous study were included in the community (JB5 - *Brevibacterium linens*, JB196 - *Vibrio casei* and JB232 - *Hafnia ssp.*)^5^. Strains of the 16-member community were inoculated at equal ratios (at 100,000 bacterial cells or fungal spores each) on 10% Cheese Curd Agar^14^. For each community and sampling conditions, triplicate communities were inoculated. Agar plates were incubated in the dark at 15℃ in a humidified plastic bag. At 24, 48, 72, and 96 hours after inoculation, plates were scrubbed with a 20%wt NaCl solution using sterilized cotton swabs in a horizontal and vertical rastering pattern, followed by a rosette (**Suppl. Fig. 12)**. On days 3, 5, 7, and 21 microbial communities were collected into 1000μL of PBS+0.05% Tween using cell scrapers; for the day 3 sample, biomass collection was done before the brine wash. Half of each sample was split for various analyses: spot plating for CFU determination, glycerol stock preparation, and DNA extraction for metagenomic short-read sequencing in the case of days 7 and 21. Half of the scraped biomass was then replated onto the same petri dish. To calculate total bacterial and fungal CFUs, spot plating of serial dilutions of each sample was done on PCAMS, either with 50μg/L chloramphenicol to count the fungal community members or 100μg/L cycloheximide + 21.6μg/L natamycin to count the bacterial community members, with colony counting done 48 hours later **(Suppl. Table 18)**. DNA was extracted from the *in vitro* community samples from days 7 and 21 using phenol-chloroform extraction (without a liquid nitrogen grinding step) and purified with 5 additional ethanol washes to remove residual phenol and chloroform. Library preparation and metagenomic sequencing of the multiplexed libraries were performed by Novogene (Sacramento, CA, USA) on a NovaSeq 6000 with a run configuration of 2 x 150 bp.

Metagenomic reads were then mapped back to the bacterial reference genomes using bwa mem to show relative abundance. For *Brevibacterium linens* (JB5, Accession number KF669529), *Vibrio casei* (JB196, Accession number Z_FUKS00000000), and *Hafnia alvei* (JB232, Accession number KF669544) previously published reference genomes were utilized. For the CCS strains the *de novo* assembled genomes from this study were utilized. 1,259,325,714 reads were sequenced, and over 99.7% of reads were aligned against the reference bacterial genomes **(Suppl. Table 20)**. Anvi’o (version 7.1 ‘hope’)^62^ was used to visualize the metagenome and the coverage from short reads. The coverage values of the contigs were filtered for nucleotide positions that were within the interquartile range (25%-75%) of coverage values for each contig before being averaged. Across the contigs of each genome, these coverage values were also averaged, weighted by their length. To calculate the relative abundance for each community member, the coverage values were divided by the sum of all coverage values. The PCA plots of the coverages were prepared in R (version 4.0.2) (https://www.r-project.org/) and the tidyverse (version 1.3.1) and ggfortify (version 0.4.14)^78^ packages. The principal component analysis was carried out using the prcomp function (center = TRUE, scale. = TRUE).

## Data availability

### Raw sequencing data

Fastq files of all metagenomic raw sequencing data are available on the Sequence Read Archive under Accession Number PRJNA778418. Subread files of the HiFi sequencing are available upon demand. Fastq files of all isolate sequencing data are available on the Sequence Read Archive under the Accession Numbers PRJNA837750 (CCS156), PRJNA837754 (CCS158), PRJNA837770 (CCS160), PRJNA837776 (CCS164), PRJNA838264 (CCS165), PRJNA837782 (CCS166), PRJNA837789 (CCS169), PRJNA838091 (CCS174), PRJNA838092 (CCS175), PRJNA838093 (CCS176), PRJNA838094 (CCS177), PRJNA838095 (CCS178), PRJNA838105 (CCS179), PRJNA838104 (CCS180), PRJNA838102 (CCS181), PRJNA838100 (CCS182), PRJNA838106 (CCS183), PRJNA838262 (CCS184), PRJNA838261 (CCS196). Fastq files of the *in vitro* community sequencing are available on the Sequence Read Archive under Accession Number PRJNA852571.

### Assemblies, MAGs, supplemental files

Assemblies, MAGs and supplemental files have been deposited on Dryad (https://doi.org/10.5061/dryad.bg79cnpd8).

## Acknowledgments

We thank Jasper Hill Farm for supplying the cheese samples, Pacific Biosciences for supplying SMRT sequencing reagents and project support as well as SNPSaurus and the Genomics & Cell Characterization Core Facility at the University of Oregon for providing SMRT sequencing services. In addition, we thank the Graduate Women in Science fellowship awarded to Christina C. Saak that allowed the generation of metaHi-C data and the IGM Genomics Center at the University of California San Diego/Illumina for a NovaSeq sequencing grant to Emily C. Pierce. We also thank Steven Villareal for help with DNA extractions from Cheese B isolates for Nanopore sequencing, Mathieu Almeida for advice on the bioinformatic analyses and Brooke Johnson for help with exploratory data analysis.

## Author contributions

C.C.S, R.J.D, M.A. and R.H. conceptualized the study. C.C.S., with help from E.C.P., collected the samples and generated the metagenomic sequencing data. C.C.S. isolated community members from Cheese B and generated their corresponding sequencing data. C.B.D. conducted the *in vitro* community experiment. C.C.S. and E.C.P. analyzed the data. D.P. generated the long-based assemblies and MAGs. C.C.S. and E.C.P. prepared the figures. C.C.S. wrote the manuscript with input from all authors. All authors agree with the contents of this manuscript.

---

**Supplemental Figure 1.**
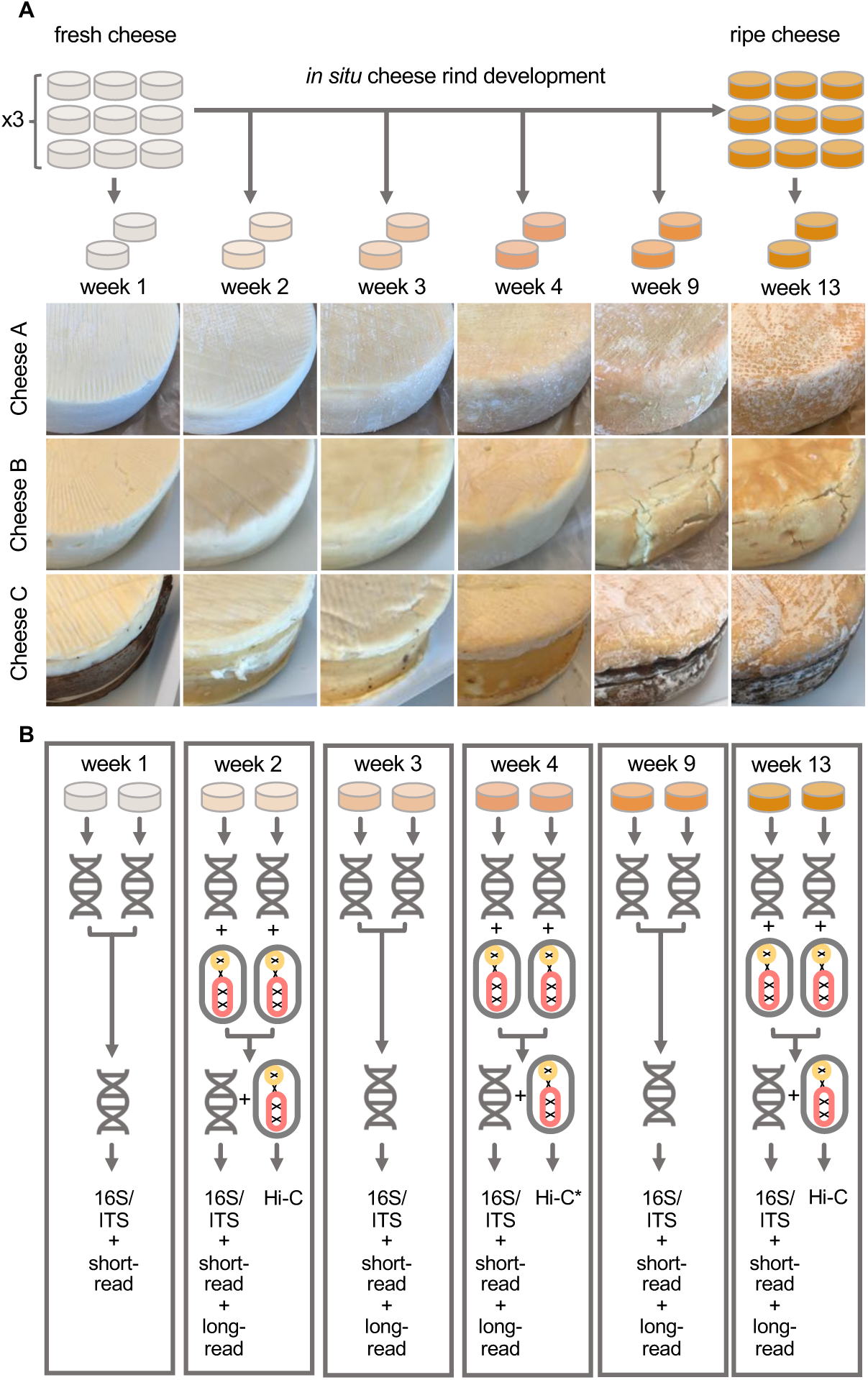
Sampling (A) and metagenomic sequencing (B) of three ripening washed-rind cheeses from the same facility. **(A)** For each of the three different washed-rind cheeses (A, B, and C) we followed the aging of three different batches produced one week apart. From each batch, we collected rind from duplicate wheels at six time points. A detailed collection schedule, including information and wash frequency and storage, can be found in **Supp. Table 1**. Representative images of each of the three cheeses at different time points are shown. Cheese C is wrapped in spruce during ripening. In the pictures for weeks 2-4 the spruce has been removed. **(B)** DNA was extracted from each rind sample collected and DNA from duplicate wheels was then combined for downstream sequencing. All three batches of each cheese were analyzed at all of the six time points with 16S and ITS amplicon sequencing to estimate reproducibility of bacterial and fungal succession dynamics, respectively. One batch (batch 3) of each cheese was analyzed at all of the six time points with short-read shotgun sequencing. The same batch of each cheese was analyzed at weeks 2, 3, 4, 9 and 13 using long-read shotgun sequencing. Finally, rind samples of that batch from weeks 2 and 13 were fixed, combined and subjected to Hi-C sequencing. *Only cheeses B and C were analyzed by Hi-C at week 4.

**Supplemental Figure 2.**
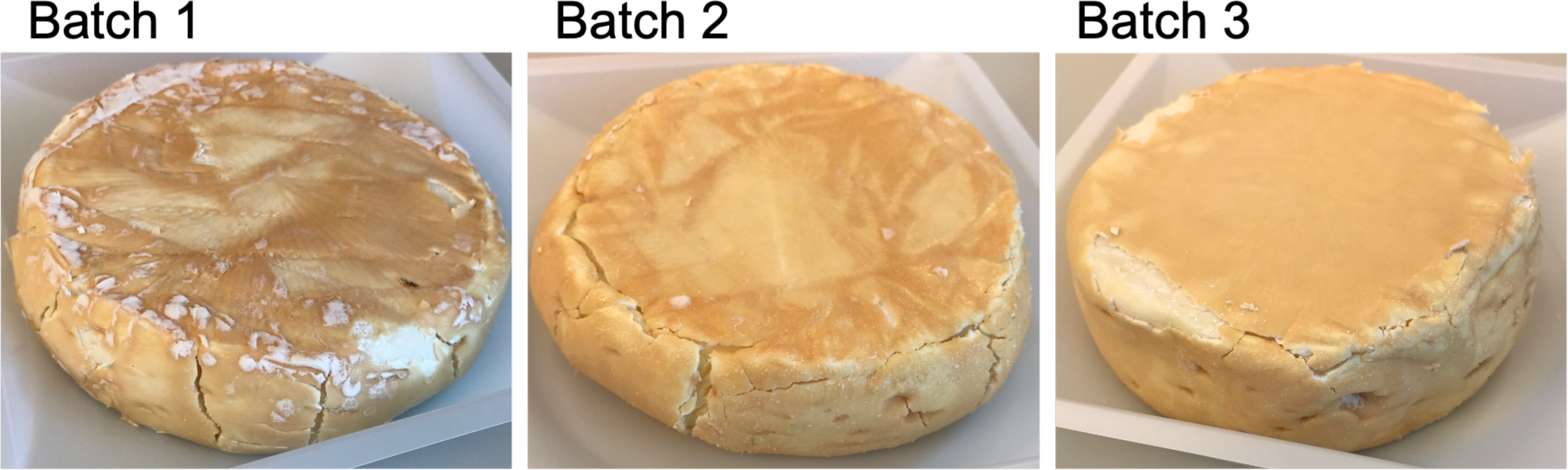
Cheese wheels from batches 1, 2 and 3 of Cheese B at week 13.

**Supplemental Figure 3.**
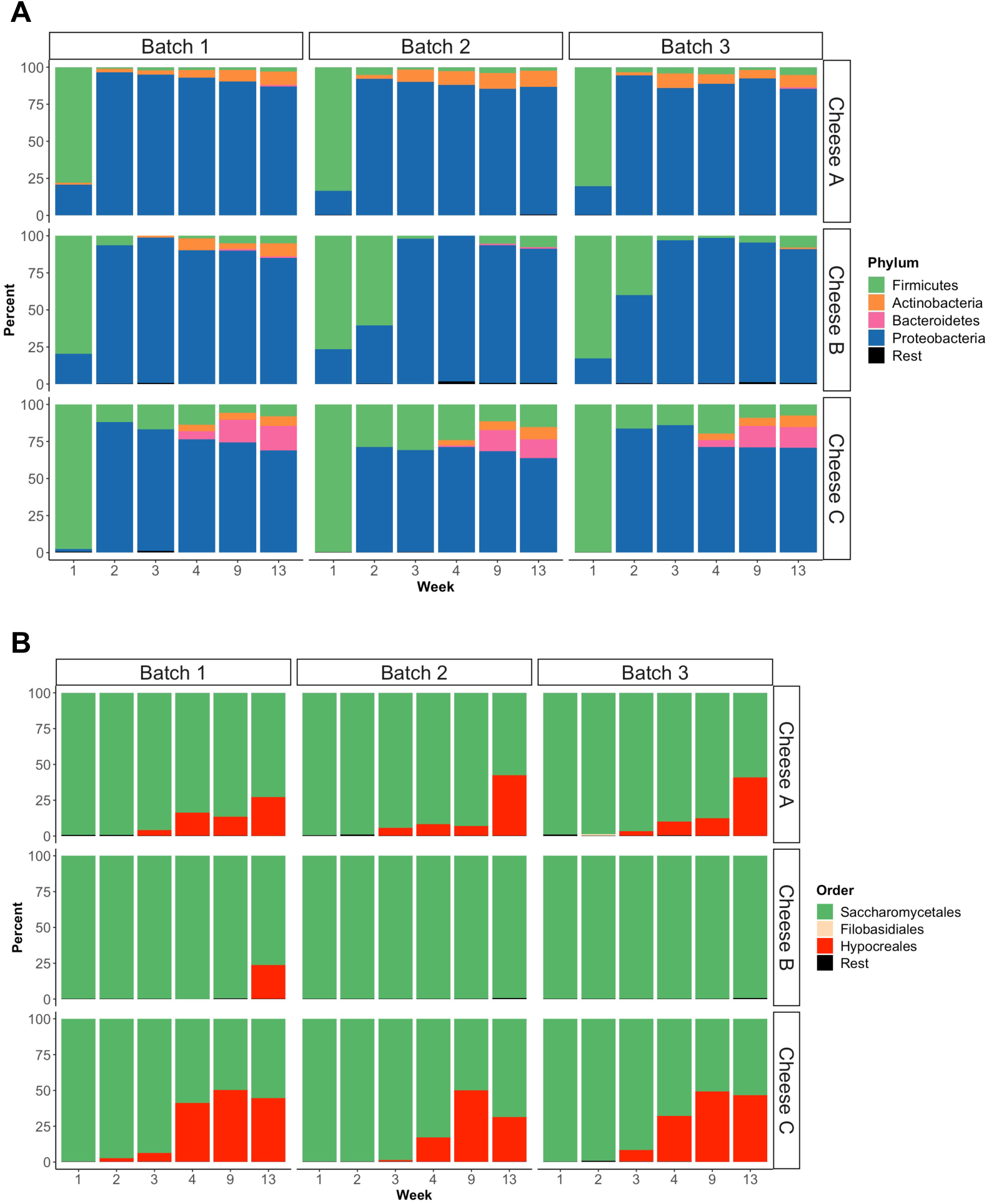
*In situ* succession patterns of bacterial and fungal washed-rind cheese communities at Phylum- and Order-level, respectively. Relative abundance plots of (A) bacteria as determined by 16S amplicon sequencing and (B) fungi as determined by ITS sequencing. Shown are the relative abundances of amplicon sequence variants collapsed at the phylum-level (A) or order-level (B). Rest = Taxa with <1% of the classified reads.

**Supplemental Figure 4.**
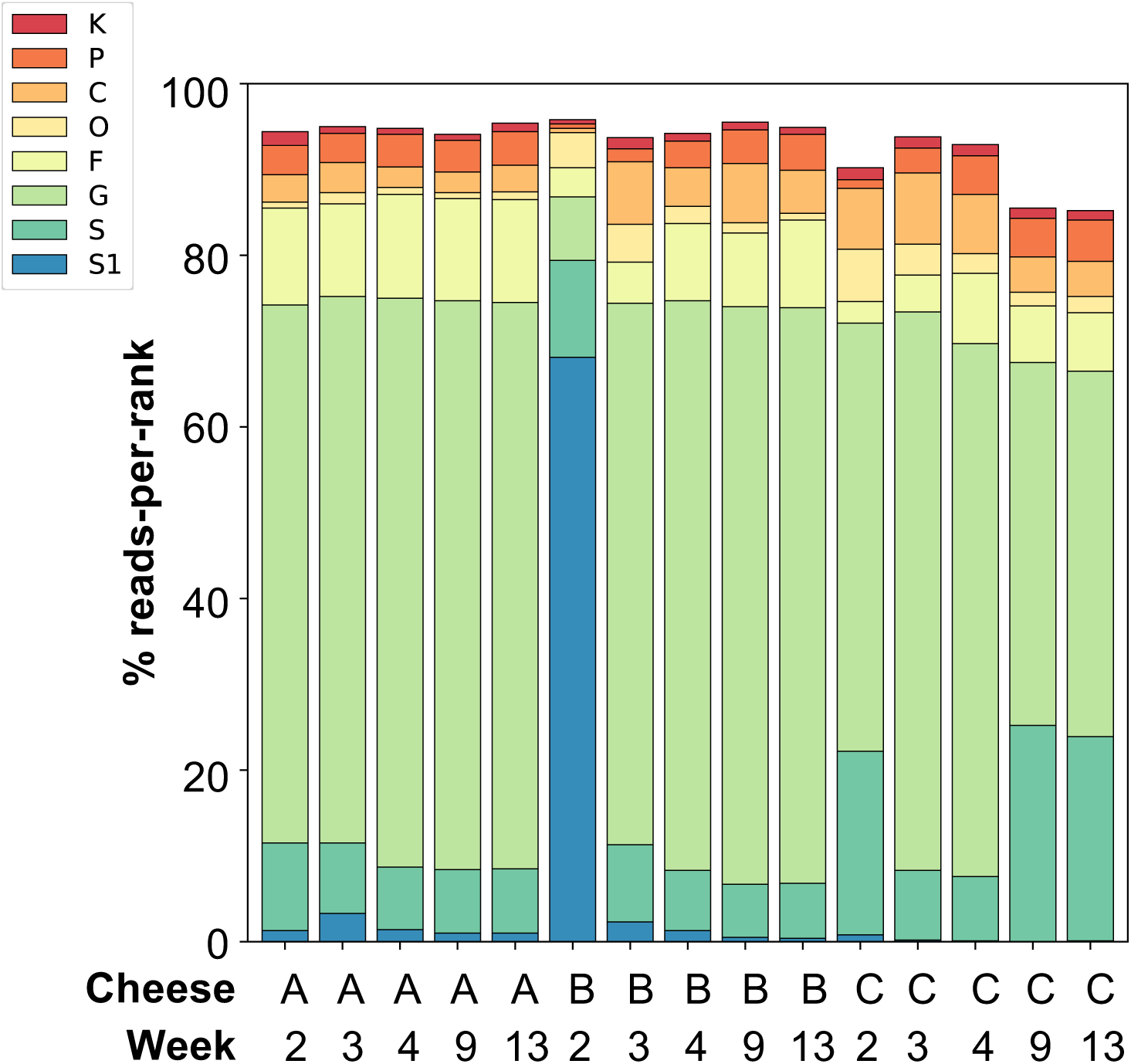
The majority of long reads were classified to at least Genus level. HiFi reads were used as inputs for the **“**Taxonomic-Profiling-Nucleotide” pipeline followed by the “MEGAN-RMA” summary. The results were visualized with the “compare-kreport-taxonomic-profiles” tool. All pipelines are part of the “PB-metagenomics-tools” toolkit (https://github.com/PacificBiosciences/pb-metagenomics-tools). Axes labels were customized in Microsoft^Ⓡ^ PowerPoint for Mac (version 16.59).

**Supplemental Figure 5.**
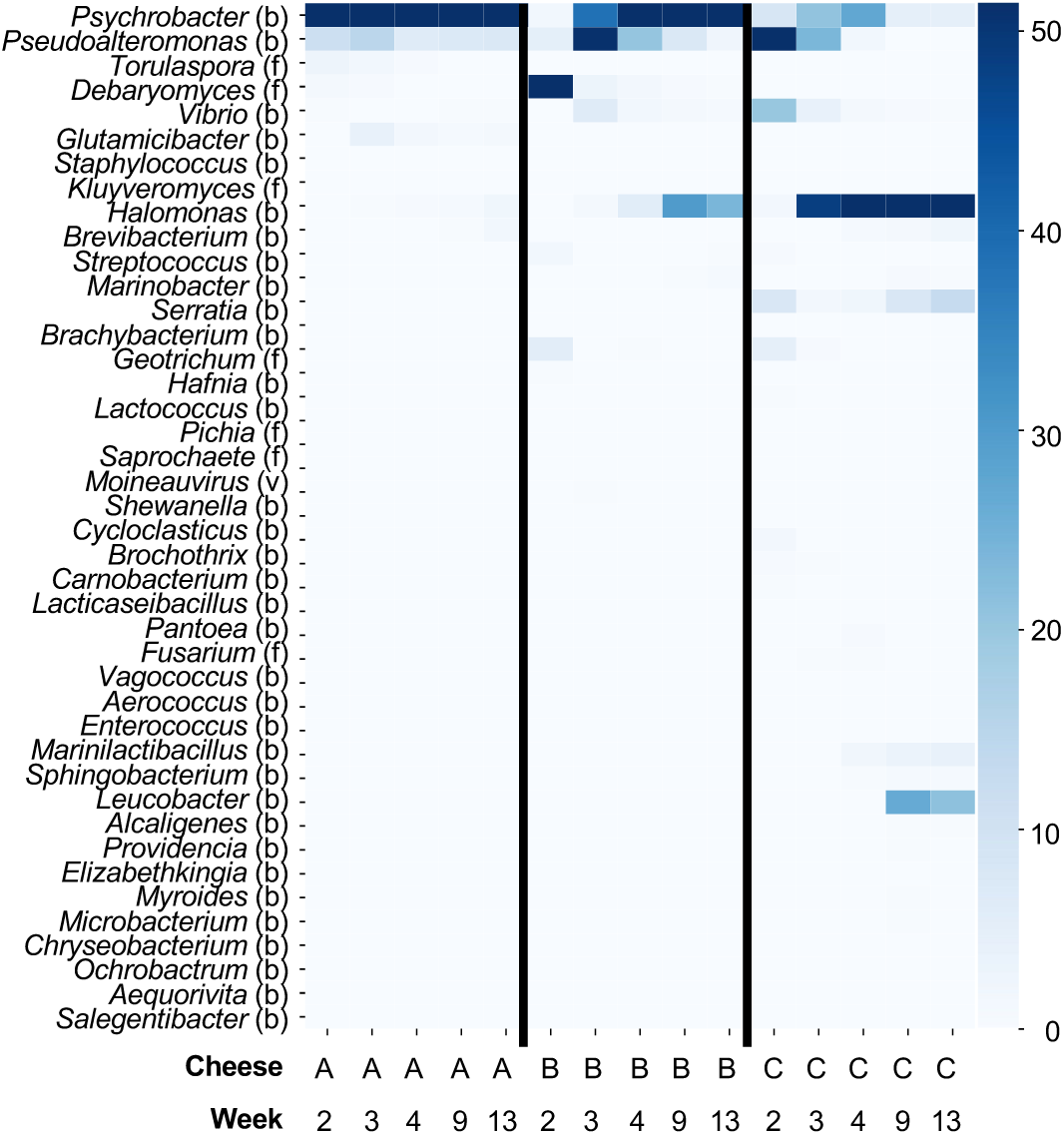
Heatmap of long-read-based relative abundance estimation aggregated at Genus level. Bacterial taxa are indicated with a (b), fungal taxa are indicated with a (f) and viral taxa are indicated with a (v). HiFi reads were used as inputs for the **“**Taxonomic-Profiling-Nucleotide” pipeline followed by the “MEGAN-RMA” summary. The results were visualized with the “compare-kreport-taxonomic-profiles” tool. All pipelines are part of the “PB-metagenomics-tools” toolkit (https://github.com/PacificBiosciences/pb-metagenomics-tools). Axes labels were customized in Microsoft^Ⓡ^ PowerPoint for Mac (version 16.59).

**Supplemental Figure 6.**
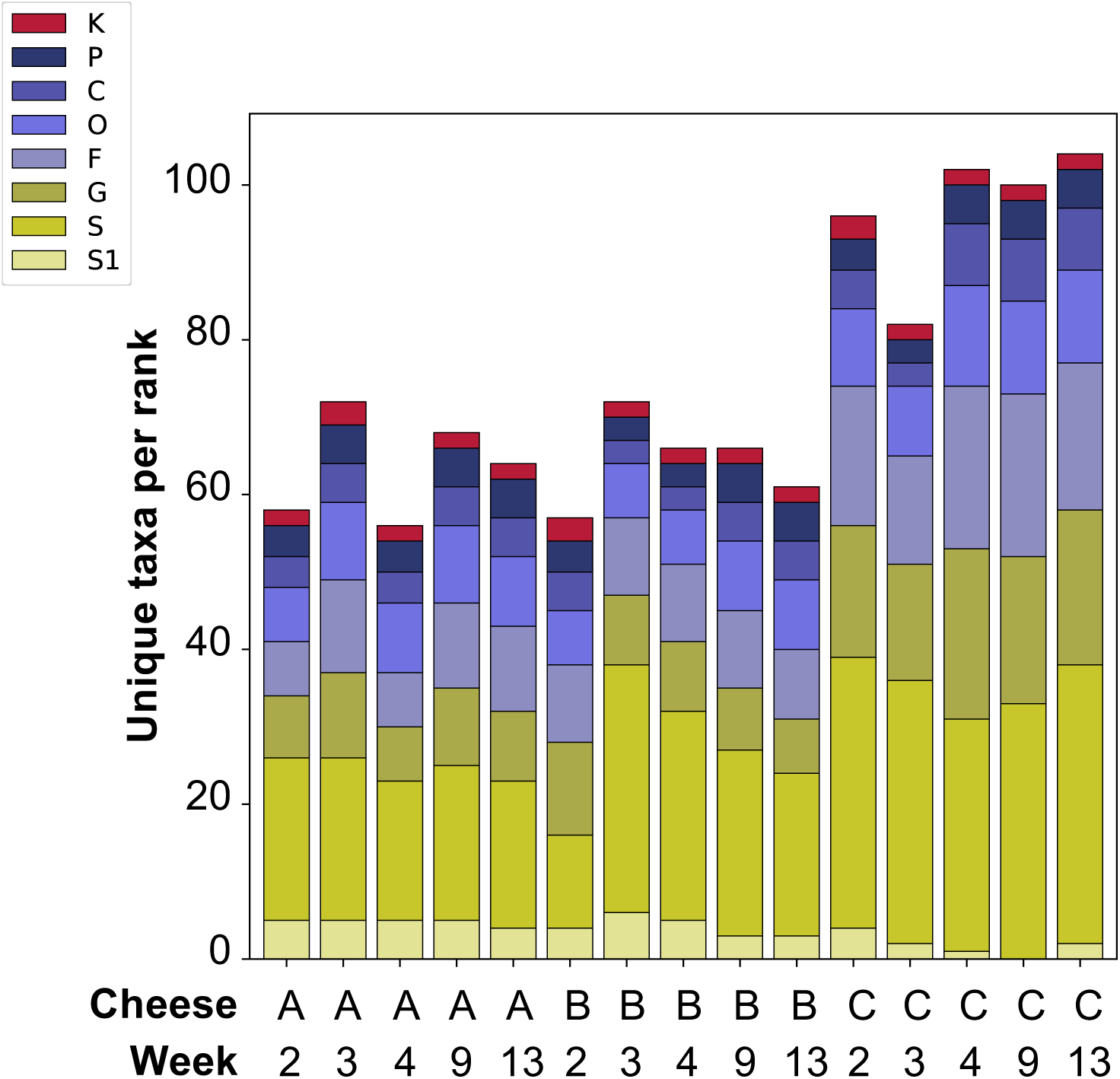
Cheese C contained the largest number of unique taxa per rank at all sampled time points. HiFi reads were used as inputs for the **“**Taxonomic-Profiling-Nucleotide” pipeline followed by the “MEGAN-RMA” summary. The results were visualized with the “compare-kreport-taxonomic-profiles” tool. All pipelines are part of the “PB-metagenomics-tools” toolkit (https://github.com/PacificBiosciences/pb-metagenomics-tools). Axes labels were customized in Microsoft^Ⓡ^ PowerPoint for Mac (version 16.59).

**Supplemental Figure 7.**
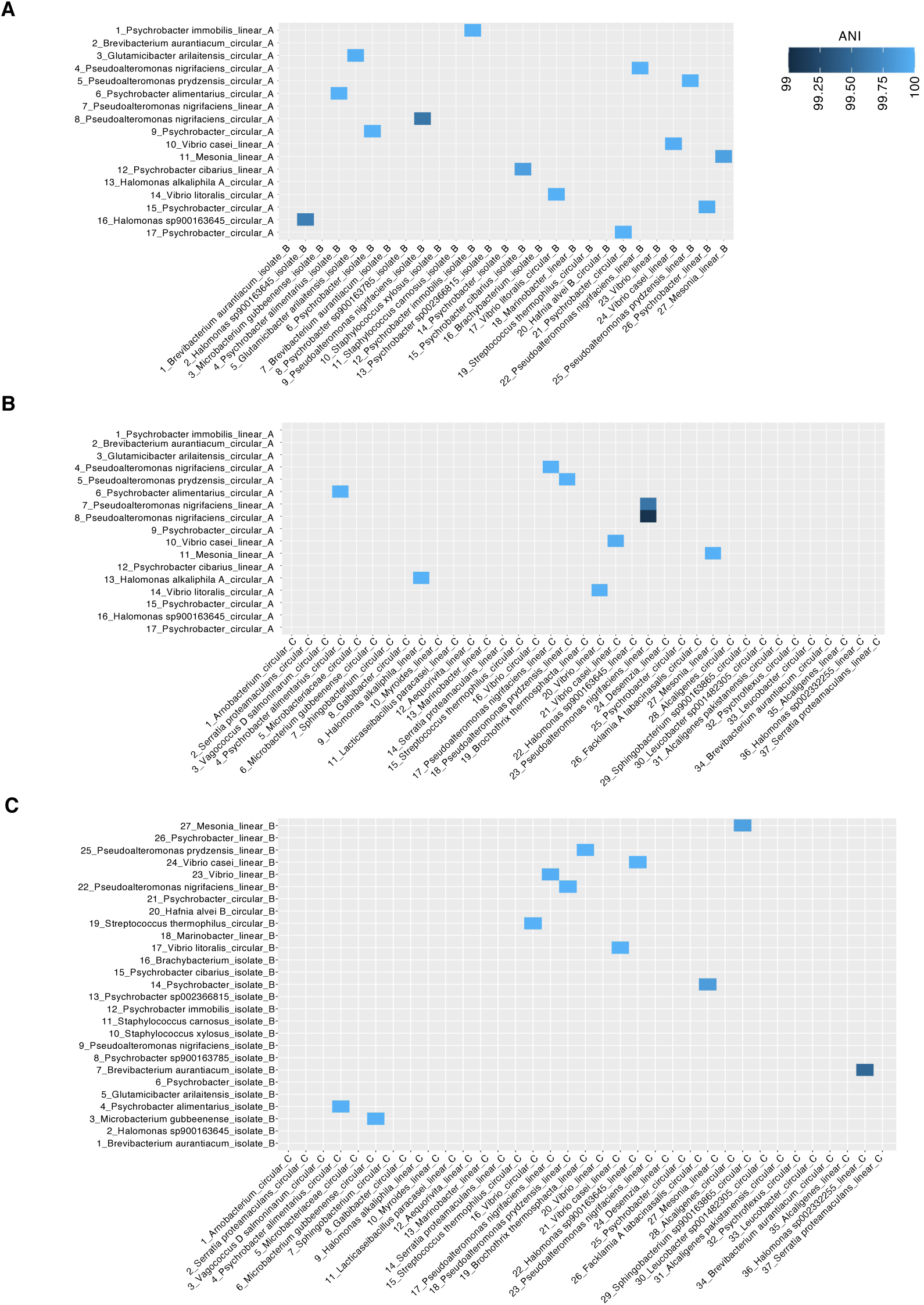
ANI Comparison of high-quality MAGs (and isolate genomes) from Cheese A, B and C. **(A)** Comparison of high-quality MAGs (and isolate genomes) recovered from cheeses A and B. **(B)** Comparison of high-quality MAGs recovered from cheeses A and C. **(C)** Comparison of high-quality MAGs (and isolate genomes) from cheeses B and C.

**Supplemental Figure 8.**
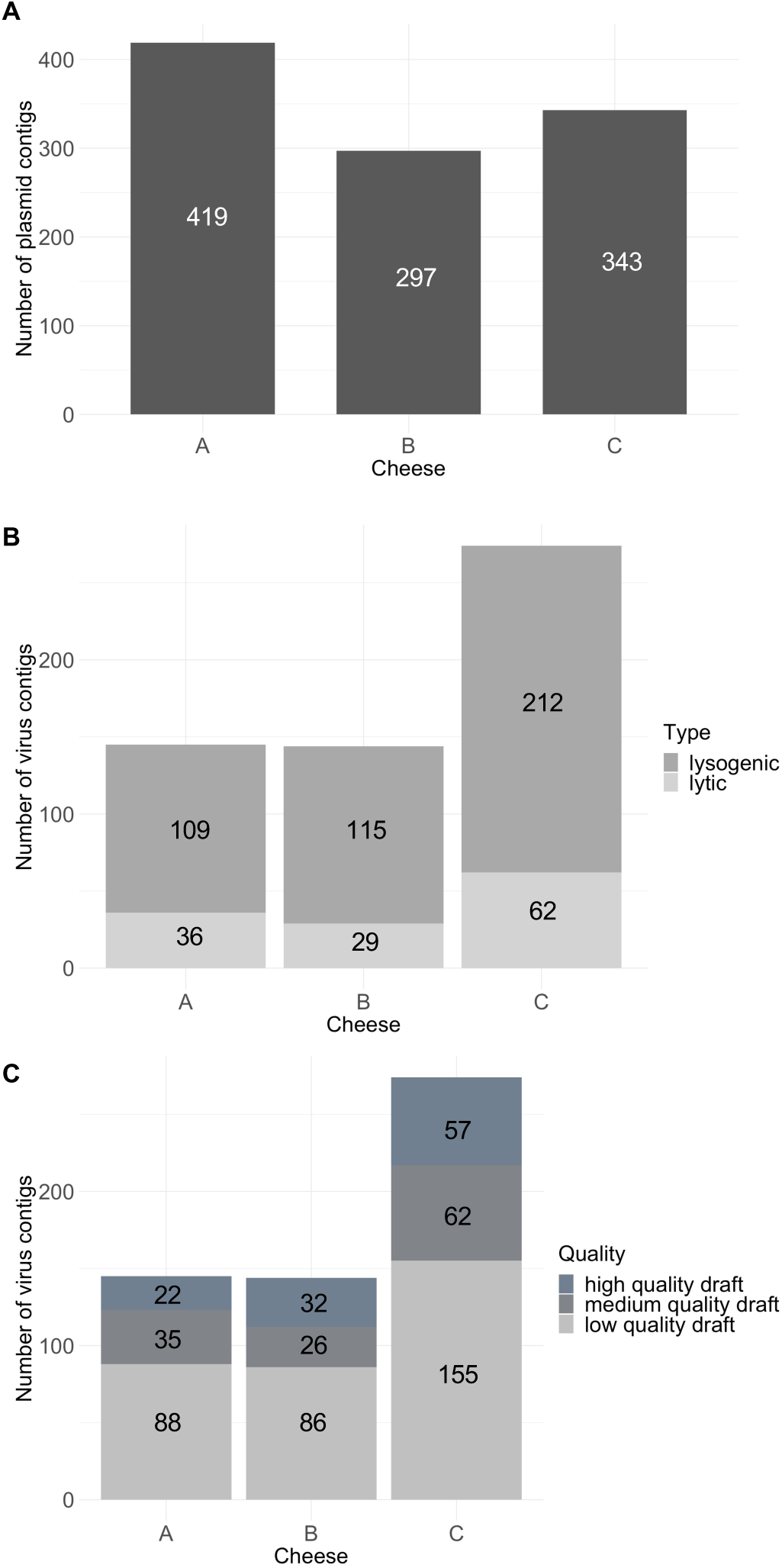
Plasmids and viruses predicted in the mega-assemblies of cheeses A, B and C. **(A)** Number of contigs predicted to be plasmids by ViralVerify (with the -p flag). **(B)** Number of contigs predicted to be lytic and lysogenic viruses by VIBRANT. **(C)** Number of high-quality, medium-quality and low-quality viral contigs as determined by VIBRANT. Of the predicted viral contigs 4, 4 and 7 are predicted to be circular for Cheeses A, B and C, respectively. Full MGE prediction results can be found in **Suppl. Table 13.**

**Supplemental Figure 9.**
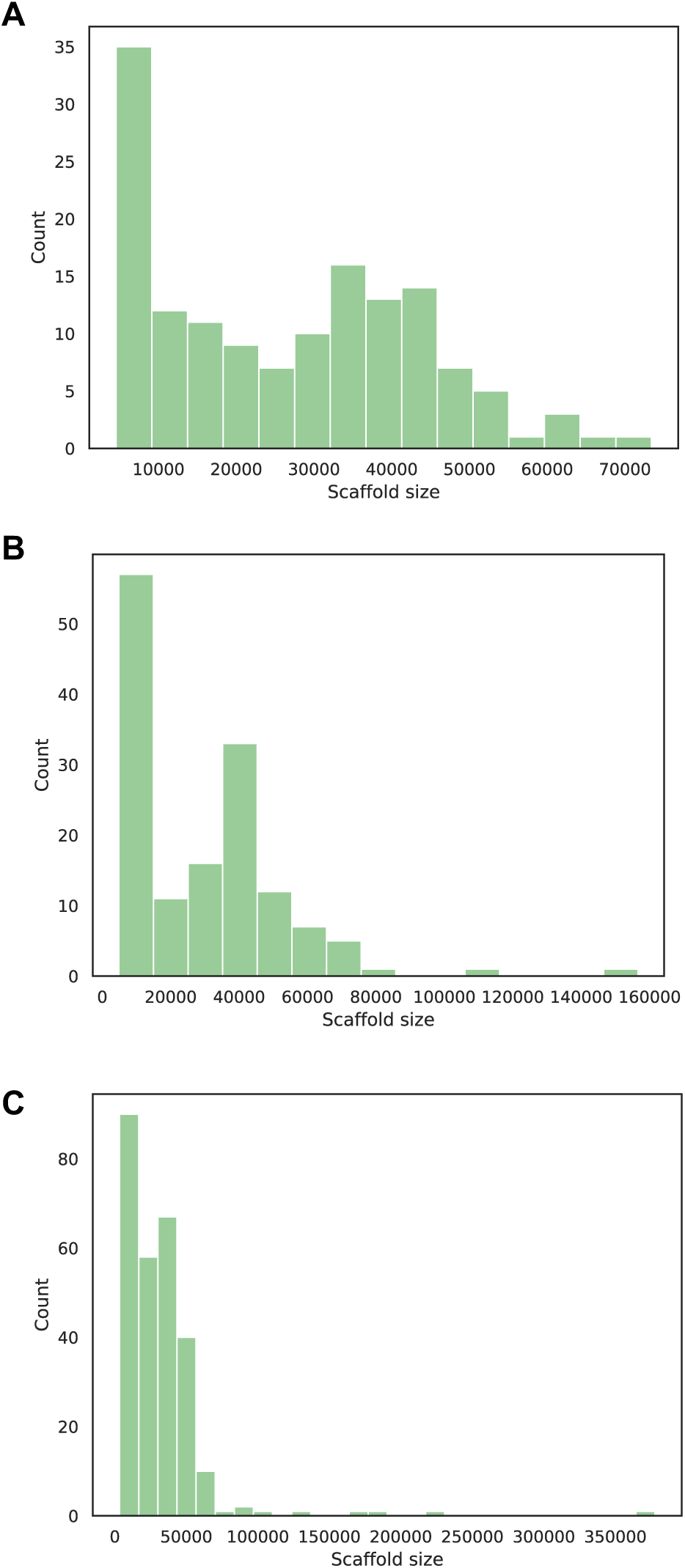
Size distributions of predicted viral contigs. Size distributions of the predicted viral contigs are shown for **(A)** cheese A, **(B)** cheese B and **(C)** cheese C. Graphs are from the standard VIBRANT output files

**Supplemental Figure 10.**
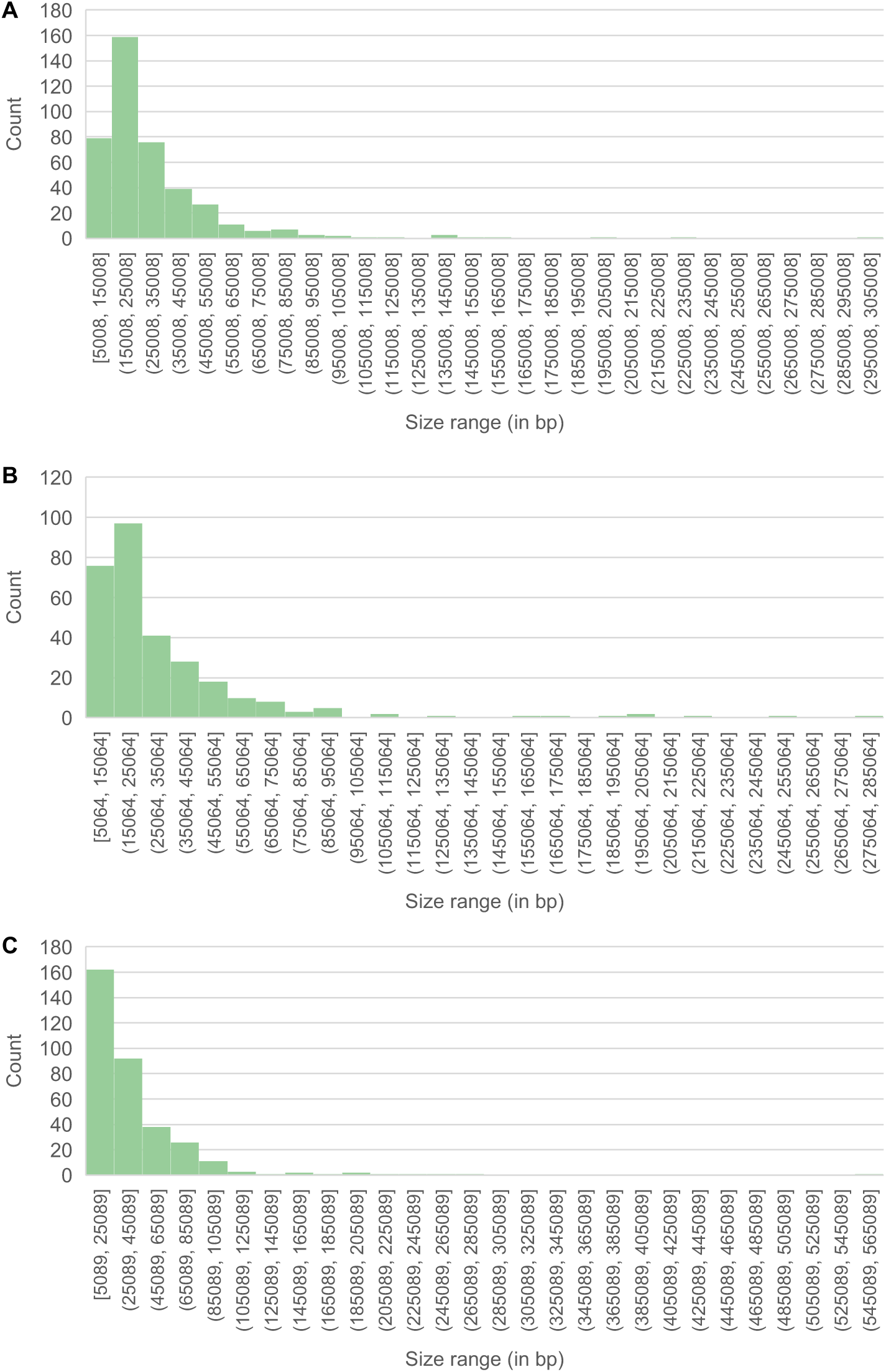
Size distributions of predicted plasmid contigs. Size distributions of the predicted plasmid contigs are shown for (A) cheese A, (B) cheese B and (C) cheese C. Graphs were produced based on ViralVerify output using Microsoft. 1206 Excel for Mac Version 16.61.

**Supplemental Figure 11.**
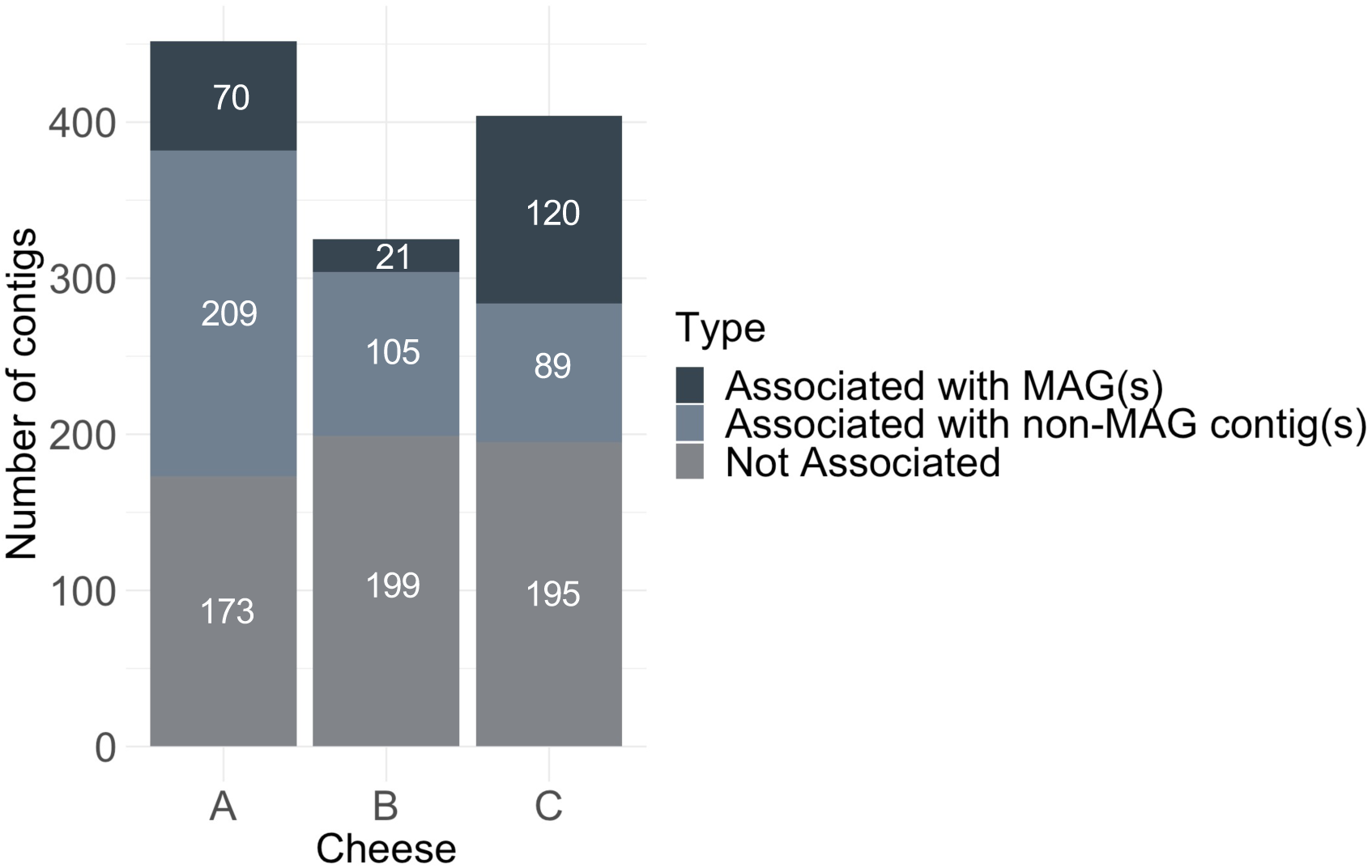
Long reads and metaHiC combined associate large numbers of lytic viruses and plasmids with hosts and unbinned contigs. For each cheese the number of lytic virus and plasmid contigs (combined due to some overlap in the classification of contigs as both lytic phages and plasmids) that is associated with MAGs or with non-MAG contigs is shown in relation to the unassociated lytic virus/plasmid contigs.

**Supplemental Figure 12.**
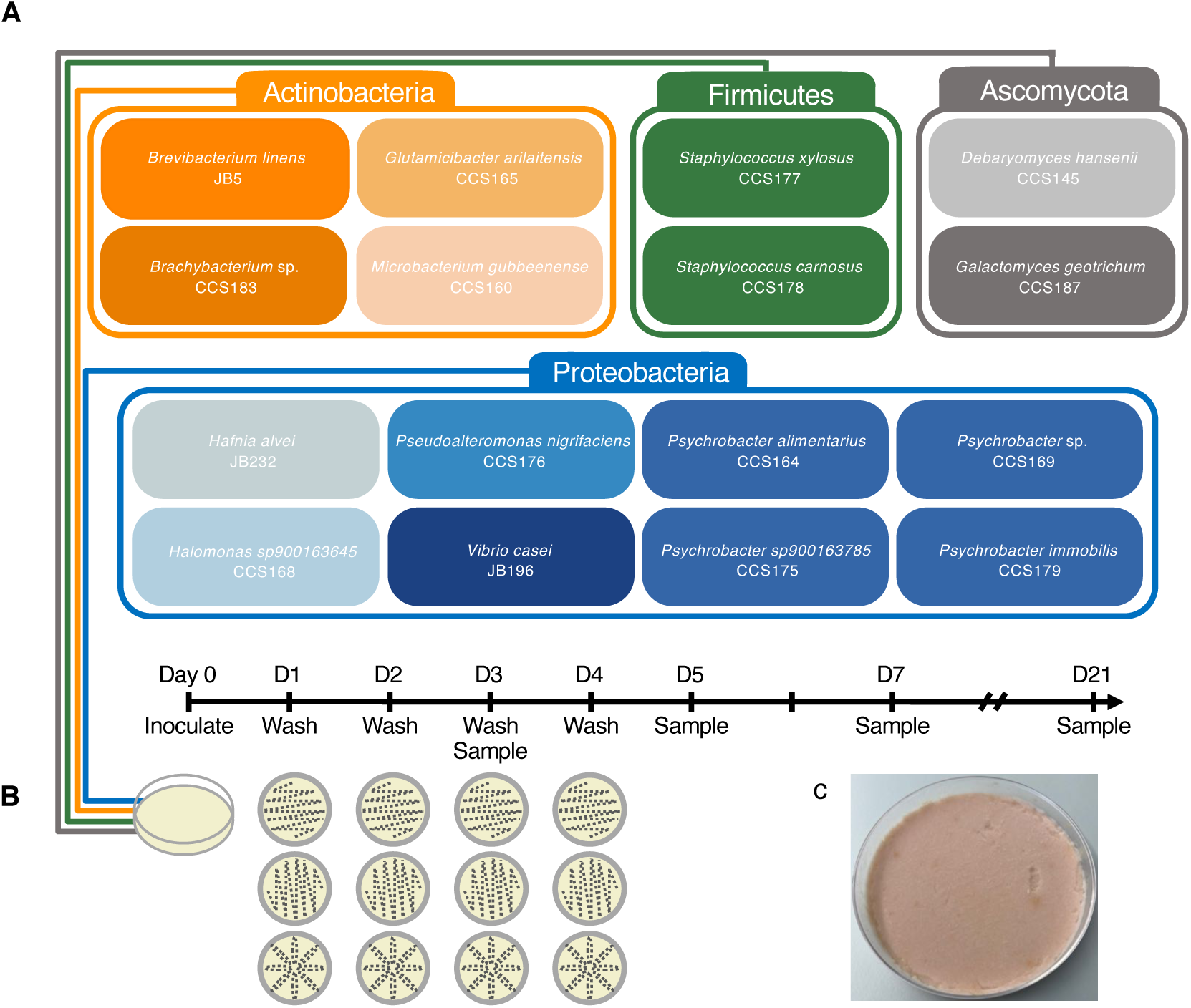
Overview of *in vitro* washed cheese rind community experiment. (a) Community members, colored by Phyla. (b) Timeline of *in vitro* microbial community model as well as washing pattern. All three steps were done on each plate with a sterile cotton swab (c) example plate of full community at day 21 from the partially destructive sampling.

**Supplemental Figure 13.**
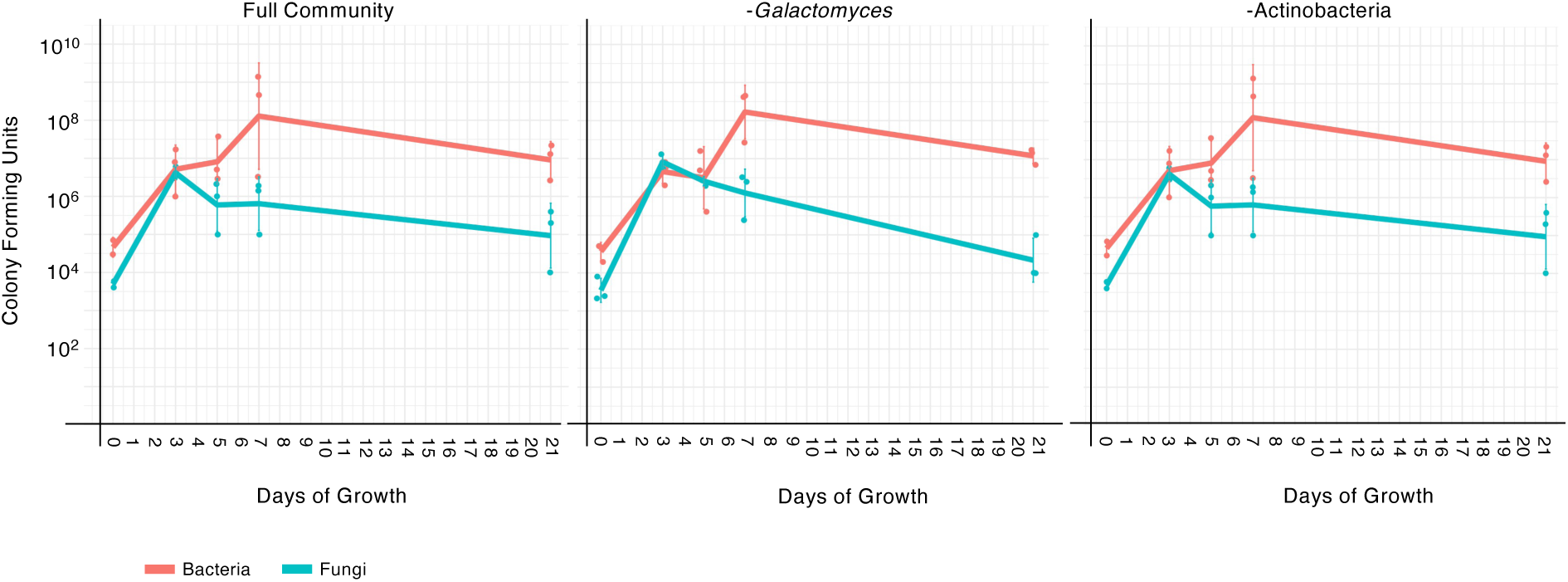
CFU counts of *in vitro* communities over time.

**Supplemental Figure 14.**
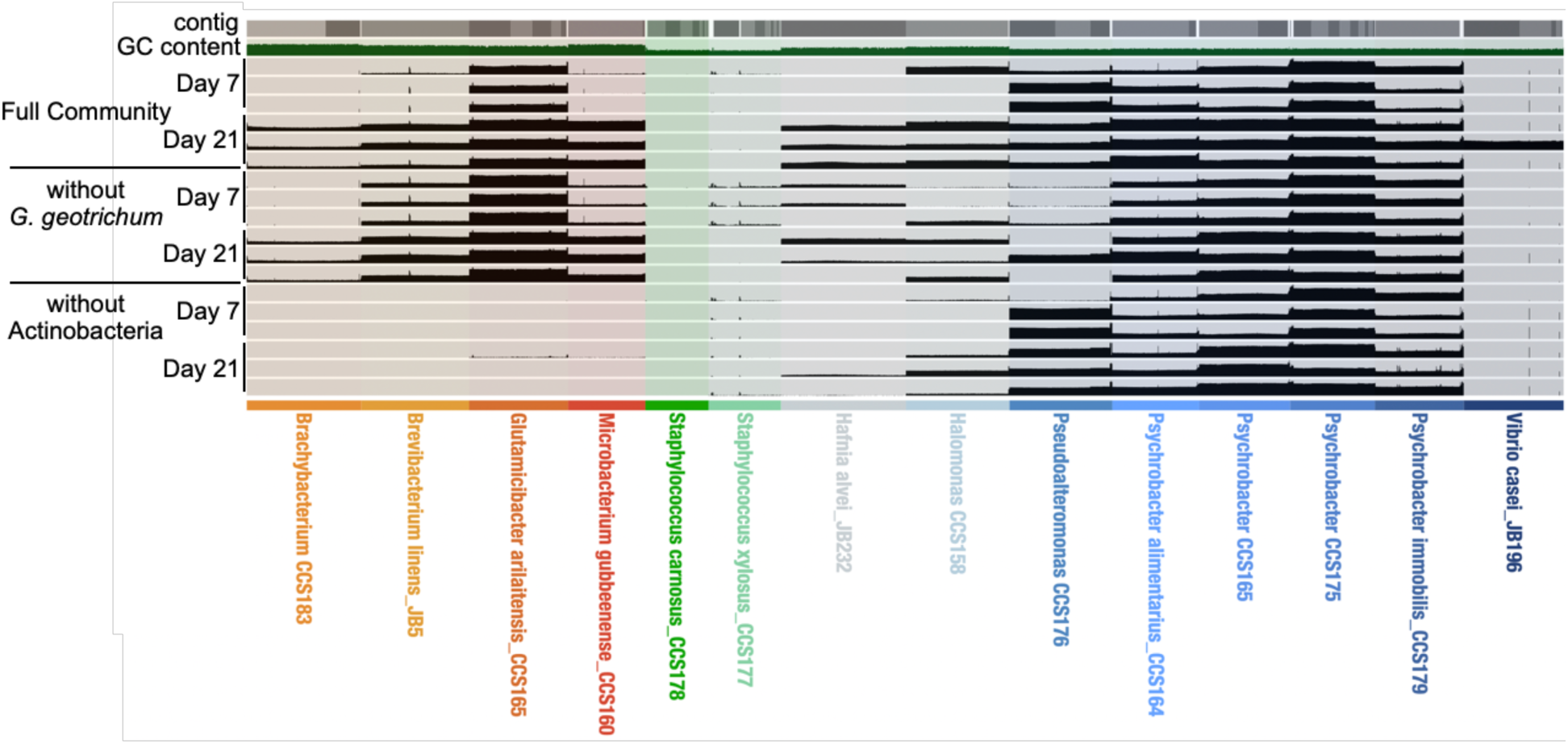
Summary of read mapping against reference bacterial genomes of *in vitro* communities over time.

**Supplemental Figure 15.**
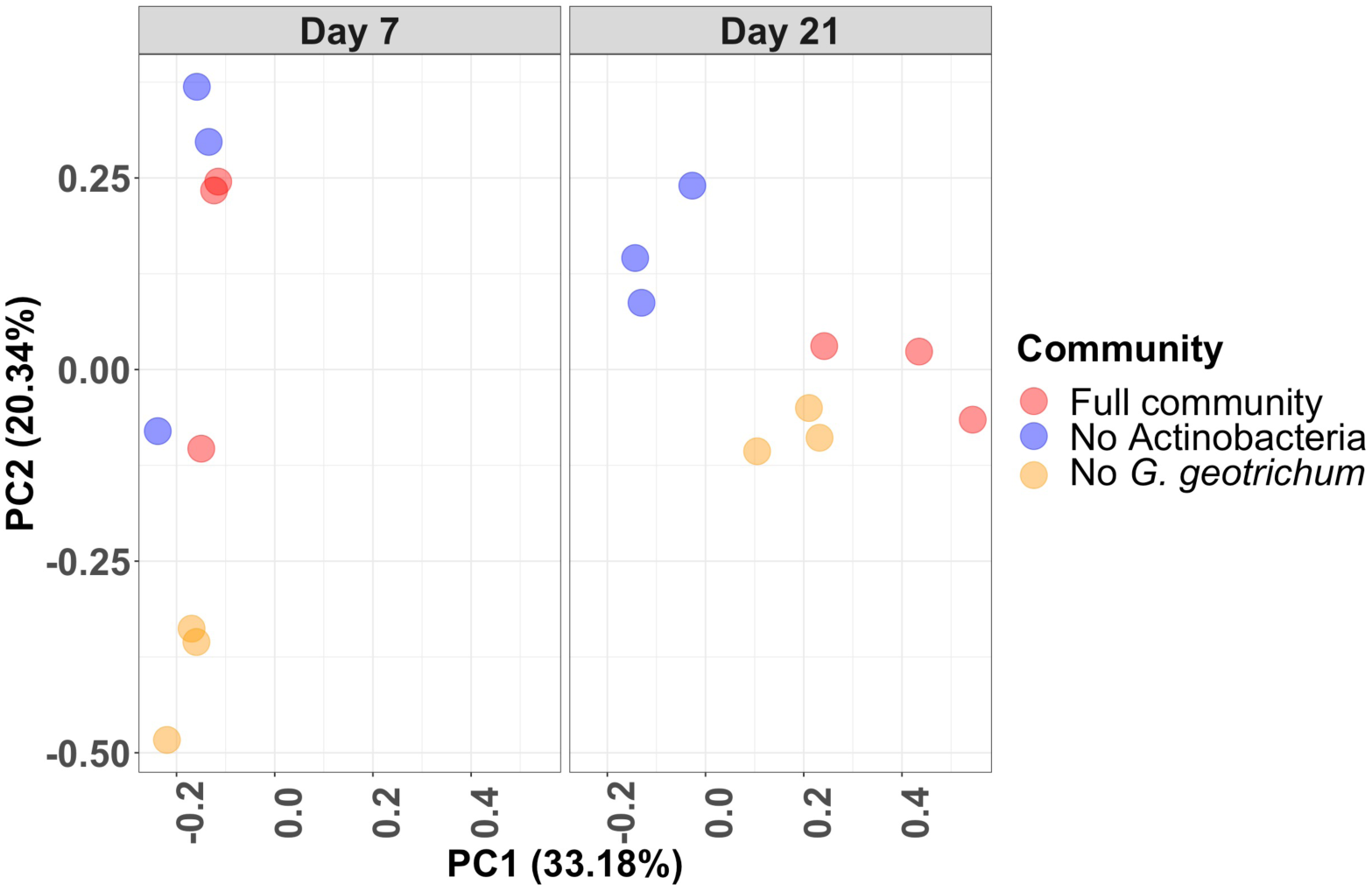
Principal component analysis of relative abundance data from *in vitro* communities.

## Supplemental Tables

**Supplemental Table 1. Collection schedule of sampled washed-rind cheeses**. For three different washed-rind cheeses (cheeses A, B and C) we followed the aging of three batches produced one week apart. Each of these batches was sampled at six time points (weeks 1, 2, 3, 4, 9 and 13). All samples were subjected to 16S and ITS amplicon (A) sequencing. All samples from batch 3 of each of the three cheeses were subjected to short-read sequencing (SR). The same set, minus the week 1 samples, were subjected to long-read SMRT sequencing (LR). A subset of batch 3 samples of each cheese was further subjected to Hi-C (H). Representative wash and storage schedules are also indicated in the columns, the exact schedules can differ from batch to batch.

**Supplemental Table 2. Sequencing data (in basepairs) generated by short-read shotgun metagenomic sequencing, long-read shotgun metagenomic sequencing and short-read Hi-C metagenomic sequencing.**

**Supplemental Table 3. 16S amplicon read classification statistics.**

**Supplemental Table 4. ITS amplicon read classification statistics. Supplemental Table 5. Long-read based taxonomic classification statistics.**

**Supplemental Table 6. Assembly statistics of individual timepoint assemblies, co-assemblies, isolates from cheese B and mega-assemblies.**

**Supplemental Table 7. Full collection of high-quality bins resulting from co- and individual timepoint assemblies.**

**Supplemental Table 8. Sequencing data (in basepairs) of short-read and long-read genome sequencing of isolates from Cheese B.**

**Supplemental Table 9. High-quality MAGs recovered from Cheese A in washed-rind cheese genomic catalog.** MAG names contain the bin number, the taxonomic classification to the lowest level available, whether it is linear or circular and which cheese it originated from. The bin name originates from the original binning procedure. Those bins that contain “Combined” in their name are based on co-assemblies, while the rest are based on individual timepoint assemblies. The number after the first letter indicates the time point they originate from (e.g. O2 originates from week 2, 2 = week 2, 3 = week 3, 4 = week 4, 5 = week 9, 6 = week 13)

**Supplemental Table 10. High-quality MAGs and isolate genomes recovered from Cheese B in washed-rind cheese genomic catalog.** MAG names contain the bin number, the taxonomic classification to the lowest level available, whether it is linear or circular or an isolate genome and which cheese it originated from. The bin name originates from the original binning procedure. Those bins that contain “Combined” in their name are based on co-assemblies, while the rest are based on individual timepoint assemblies. The number after the first letter indicates the time point they originate from (e.g. Wy2 originates from week 2, 2 = week 2, 3 = week 3, 4 = week 4, 5 = week 9, 6 = week 13)

**Supplemental Table 11. High-quality MAGs recovered from Cheese C in washed-rind cheese genomic catalog.** MAG names contain the bin number, the taxonomic classification to the lowest level available, whether it is linear or circular and which cheese it originated from. The bin name originates from the original binning procedure. Those bins that contain “Combined” in their name are based on co-assemblies, while the rest are based on individual timepoint assemblies. The number after the first letter indicates the time point they originate from (e.g. We2 originates from week 2, 2 = week 2, 3 = week 3, 4 = week 4, 5 = week 9, 6 = week 13)

**Supplemental Table 12. ANI Comparison of high-quality MAGs across the three cheeses.**

**Supplemental Table 13. Percentage of Illumina short reads that mapped to the genomic catalog.**

**Supplemental Table 14. Plasmid and virus predictions of mega-assemblies.** Contigs predicted as “complete circular” by VIBRANT are listed twice, once to indicate circularity and once to indicate predicted quality.

**Supplemental Table 15. Full results from the viralAssociationPipeline.** Each tab contains results from a single timepoint of a single cheese. Rows highlighted in yellow indicate that the host contig of the MGE was one of the high-quality MAGs recovered in this study.

**Supplemental Table 16. Potential host changes of MGEs over time.** For each cheese, we identified MGEs that were associated with at least one new host genus (as predicted by Kraken taxonomy and the viralAssociationPipeline) at a later timepoint relative to any previous time points, indicating a potential host change. Rows highlighted in yellow indicate that the host contig of the MGE at that timepoint was one of the high-quality MAGs.

**Supplemental Table 17. *Psychrobacter* genomes used for pangenome analysis.** Database identifier and isolation source for each of the genomes is provided.

**Supplemental Table 18. Functional enrichment in *Psychrobacter* genomes from cheese versus other environments.** Functional categories used in Figure 5C for categories with an adjusted q-value <0.1 are listed in the far right column.

**Supplemental Table 19. CFU counts of *in vitro* washed rind communities.**

**Supplemental Table 20. Isolate genome statistics and coverage at days 7 and 21.**

**Supplemental Table 21. “Cloud” gene clusters unique to individual *Psychrobacter* genomes from Cheese B.**

## Notes

https://doi.org/10.5061/dryad.bg79cnpd8

### Competing Interest Statement

The authors have declared no competing interest.

